# Chromosomal evolution and polyploidy dynamics in tribe Andropogoneae (Poaceae: Panicoideae): insights from phylogenomic reconstruction

**DOI:** 10.64898/2025.12.09.693223

**Authors:** Ana Tereza Hesse Bispo, Christian Silva, Cassiano A. Dorneles Welker

## Abstract

- The dissertation explored the chromosomal evolution of 230 species in the Andropogoneae tribe (Poaceae), which includes 1.224 species, using a detailed search of literature and databases. The phylogenetic analysis was based on complete plastome sequencing, and ancestral reconstruction utilized Bayesian inference and character optimization.
- Results indicated high diversity, with 46 variations in chromosome number (*2n*), ranging from 8 to 140, and 30% of species exhibiting polyploidy. This suggests multiple chromosomal alteration events occurred during the tribe’s evolution.
- The chromosome number *2n* = 20 was identified as the ancestral state for most internal nodes of the tree and confirmed as predominant in the Andropogoninae subtribe. However, more recently divergent taxa displayed higher chromosome numbers.
- The research concludes that chromosomal variation is dynamic and complex. It emphasizes the importance of obtaining more chromosomal data to fill knowledge gaps and achieve a more complete understanding of the diversity and evolutionary mechanisms shaping the Andropogoneae tribe.

## Introduction

The Poaceae family (Gramineae) is recognized as one of the largest among Angiosperms, harboring approximately 11,783 species distributed across 12 subfamilies, 54 tribes, and 789 genera, according to the current classification (Soreng et al., 2022). Its economic relevance is undeniable, as it includes vital crops for human consumption, such as sugarcane (*Saccharum officinarum L.*), maize (*Zea mays L.*), wheat (*Triticum aestivum L.*), and rice (*Oryza sativa L.*), in addition to being essential for animal feed and biofuel production (Rúgolo de Agrasar & Puglia, 2004; Judd et al., 2009). The phylogenetic structure of the family features three basal lineages and two major clades: the BOP (Bambusoideae, Oryzoideae, and Pooideae) and the PACMAD, which includes the monophyletic Panicoideae subfamily.

The Panicoideae subfamily is monophyletic and extremely diverse, comprising 3.325 species distributed across 242 genera (Soreng et al., 2022). Within it, the Andropogoneae tribe stands out as a significant monophyletic group, encompassing 1.224 species in 14 subtribes and 92 genera (Welker et al., 2020). This tribe has a broad pantropical distribution in grassland areas (Sanchez-Ken & Clark, 2010; Kellogg, 2015) and is characterized by individuals that, for the most part, have spikelets arranged in pairs (Kellogg, 2000). The genome of species within this order tends to differentiate and duplicate more when compared to other monocot groups (Leitch & Leitch, 2010), with the basic chromosome number for Andropogoneae species being x = 10 (Spangler et al., 1999).

The investigation of evolution within this complex group is carried out by plant cytotaxonomy, which integrates cytology and taxonomy, and considers the chromosome number to be the most fundamental data point, as it is not influenced by environmental factors or the plant’s developmental stage (Raven, 1975; Guerra, 2008). Changes in genome size and chromosome number are crucial evolutionary mechanisms linked to speciation and environmental adaptations (Elliott et al., 2022). Polyploidy (the presence of two or more genomes), which affects about 35% of vascular plants, is a central event in the evolution of wild and cultivated species, conferring adaptive advantages such as greater resistance and larger organs (Stebbins, 1971; Peer et al., 2020; Levin, 2002). Allopolyploidy is particularly common in the Andropogoneae tribe (Estep et al., 2014). Another relevant mechanism is dysploidy, involving structural rearrangements like centric fusion and fission, which alter the chromosome number and aid in recognizing the ancestral basic number of the studied group (Stebbins, 1971; Guerra, 1990).

Given the significant ecological importance and economic impact of Andropogoneae species, and due to the scarcity of detailed analyses on their chromosomal evolution, the present study aimed to compile and analyze the available data in the literature on the chromosome numbers of the tribe. The specific objectives were focused on reviewing the chromosomal evolution, analyzing the observed diversity, and identifying gaps in the scientific knowledge related to the species in this group.

## Materials and Methods

### Sampling and Chromosome Data Survey

The work started with a thorough search of scientific papers to gather everything available on chromosome numbers in species from the Andropogoneae tribe. Besides published articles, the research also used data from online databases like the *Chromosome Counts Database* and the *Index to Plant Chromosome Numbers*. In total, chromosome information for 230 species of Andropogoneae, along with some from closely related tribes, was collected using the phylogenetic tree by Welker et al. (2020) as a reference. All the data found, including diploid numbers (*2n*) and any synonyms of species names, were organized in spreadsheets by genus and subtribe, creating a solid information base for the later analyses.

### Data Analysis

For the analysis of chromosomal evolution, the Andropogoneae phylogenetic tree developed by Welker et al. (2020), which is based on complete plastome sequencing, was utilized. The reconstruction of the ancestral chromosome number for each node of the phylogeny was performed following the methodology proposed by Angulo et al. (2022). The collected diploid chromosome numbers (*2n*) were transformed into character states and subsequently combined with the phylogenetic tree. Species included in the Welker et al. (2020) study for which chromosomal data were not available in the literature or databases were, nonetheless, maintained in the phylogenetic tree reconstruction.

The initial data processing involved the *Geneious software* for visualization and editing of plastome sequencing, as well as for sequence alignment and the generation of the matrix in Nexus format. The phylogenetic inference stage was preceded by the selection of the most appropriate evolutionary model, which was determined by the MrModeltest software based on statistical criteria, resulting in the choice of the GTR+G+I model. Bayesian Inference was selected as the method for phylogenetic tree reconstruction, using the following parameters for the Monte Carlo Markov Chain (MCMC): 100.000.000 generations, two independent runs, and saving one phylogenetic tree every $1,000$ generations. This analysis was executed on the *Cyber Infrastructure for Phylogenetic Research* (CIPRES) platform, which processed the matrix using the MrBayes 3.2.7a tool to generate the consensus phylogenetic tree.

Finally, the consensus phylogenetic tree obtained from CIPRES and the table containing the chromosome data and their states were loaded into the *Mesquite software*. Phylogenetic optimization was performed in this environment using the principle of Maximum Parsimony, which aims to identify the tree requiring the fewest evolutionary changes to explain the distribution of chromosomal characters among the analyzed species.

## Results

### Chromosome Data

The bibliographic survey and database searches provided the diploid chromosome numbers (*2n*) for the selected species, as detailed in Table 1. For the species belonging to the Andropogoneae tribe, 46 variations in the *2n* chromosome number were found, covering the range from *2n* = 8 to *2n* = 140. A notable finding is that 69 Andropogoneae species (approximately 30% of the total sample) exhibited more than one reported chromosome number. For the phylogenetically close species, 11 variations in the *2n* chromosome number were identified (Table 2).

**Table 1.**
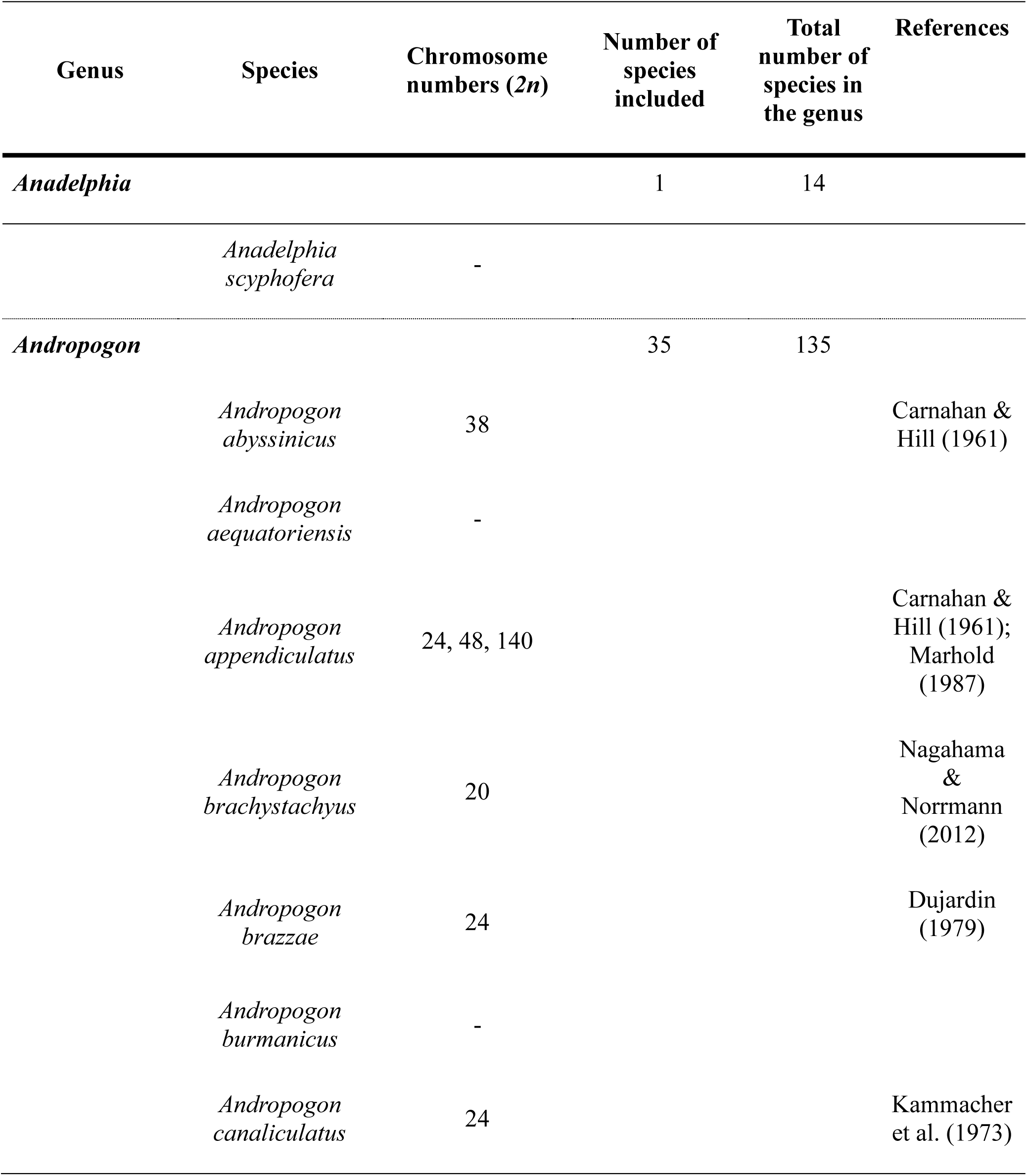
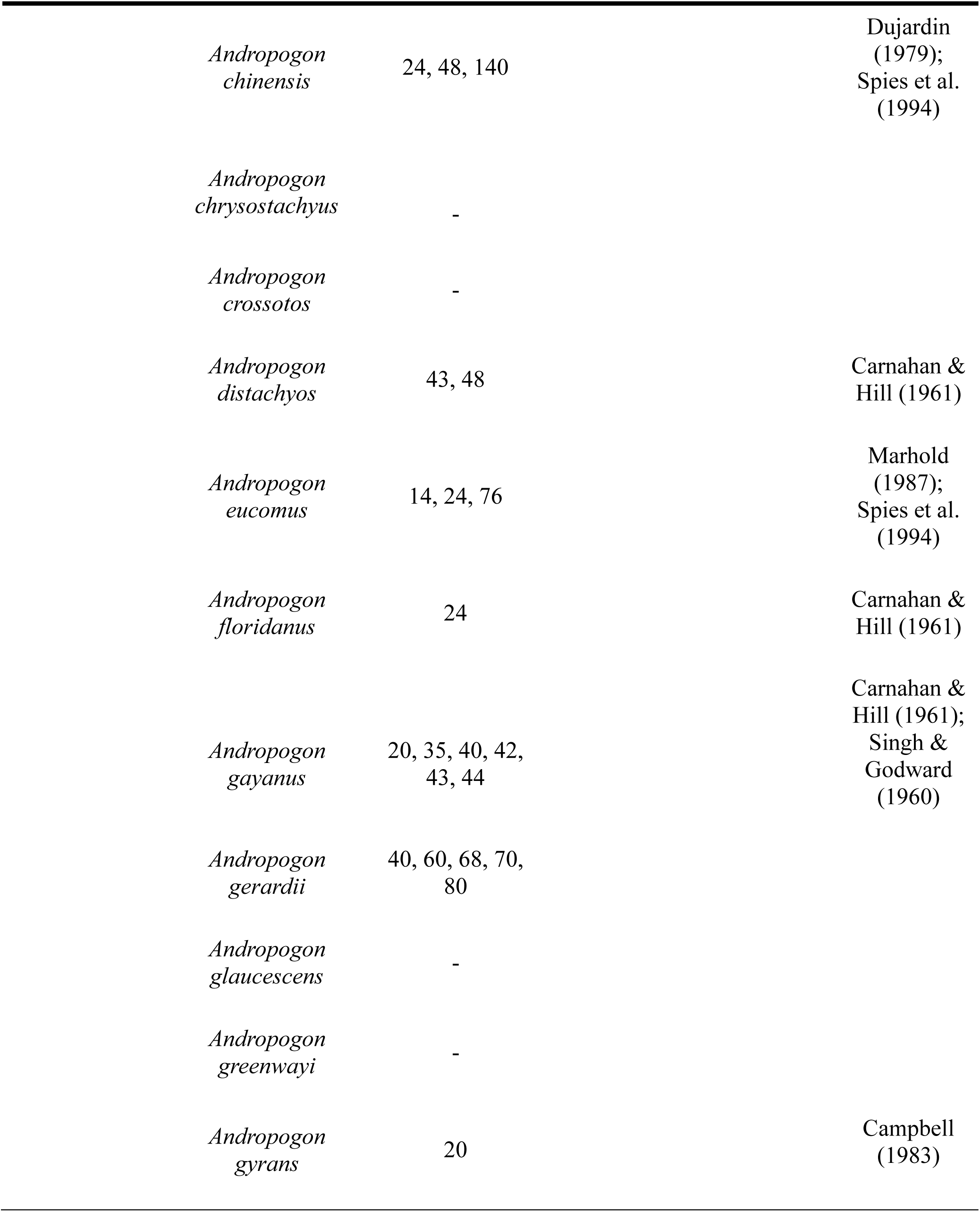
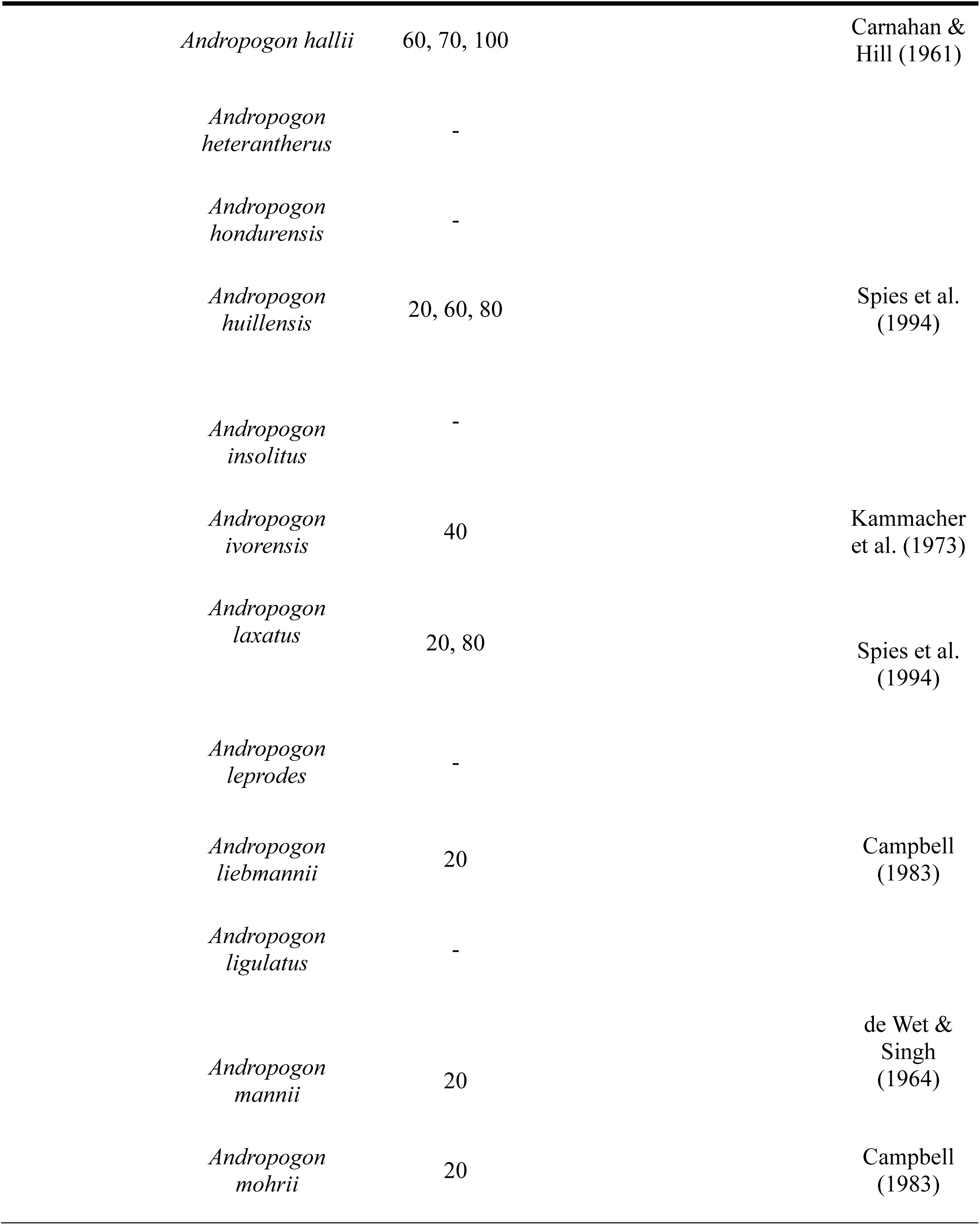
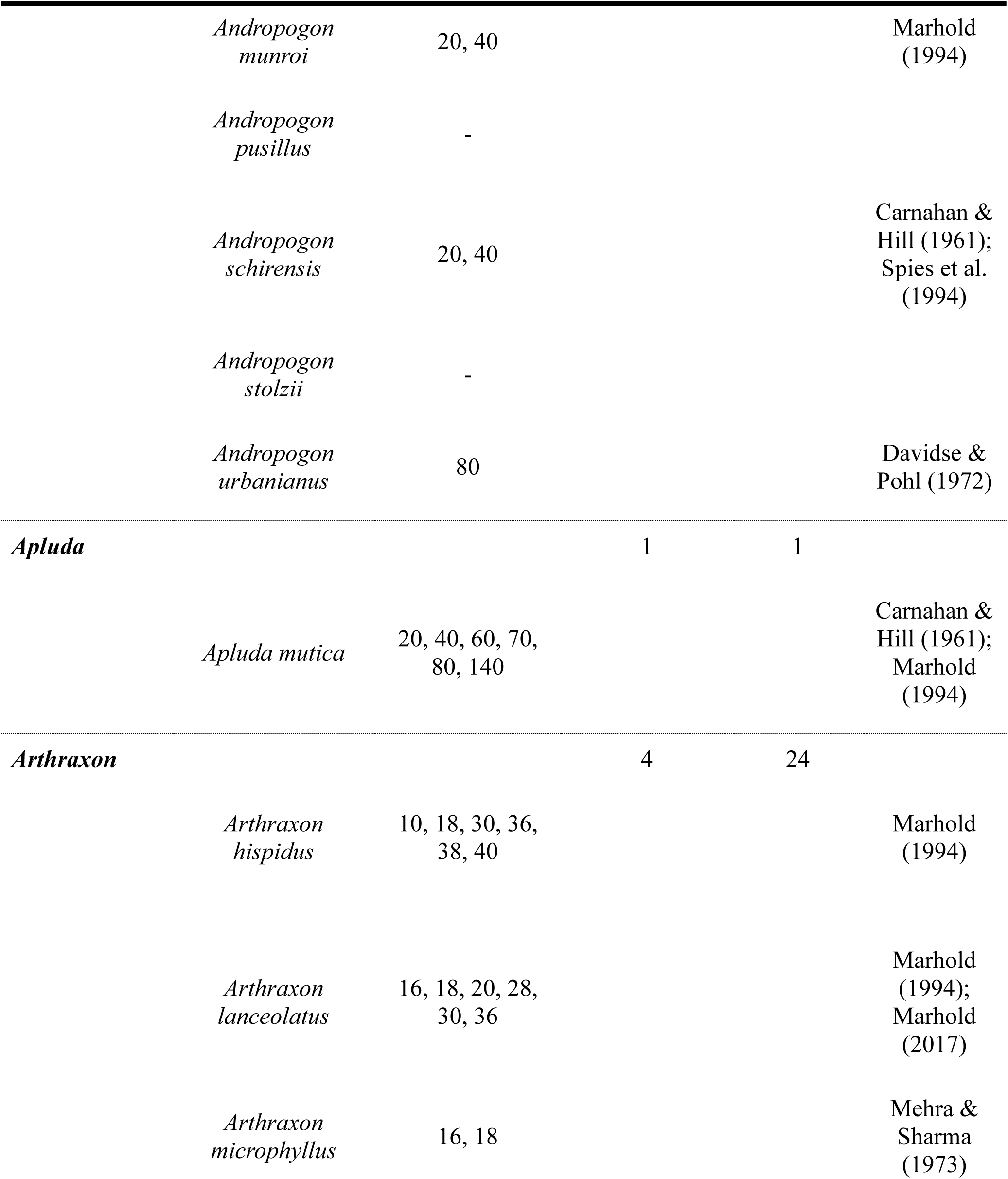
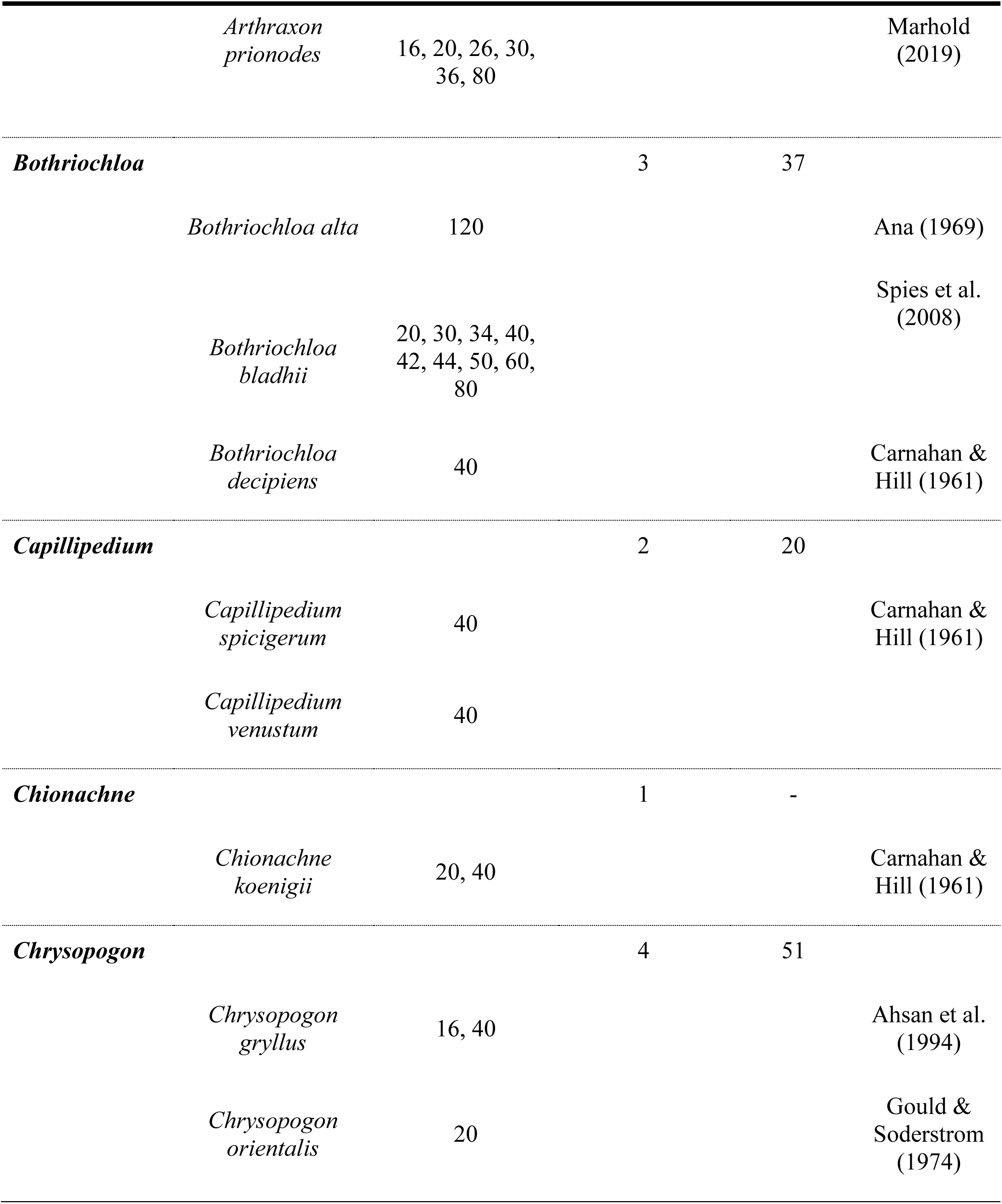
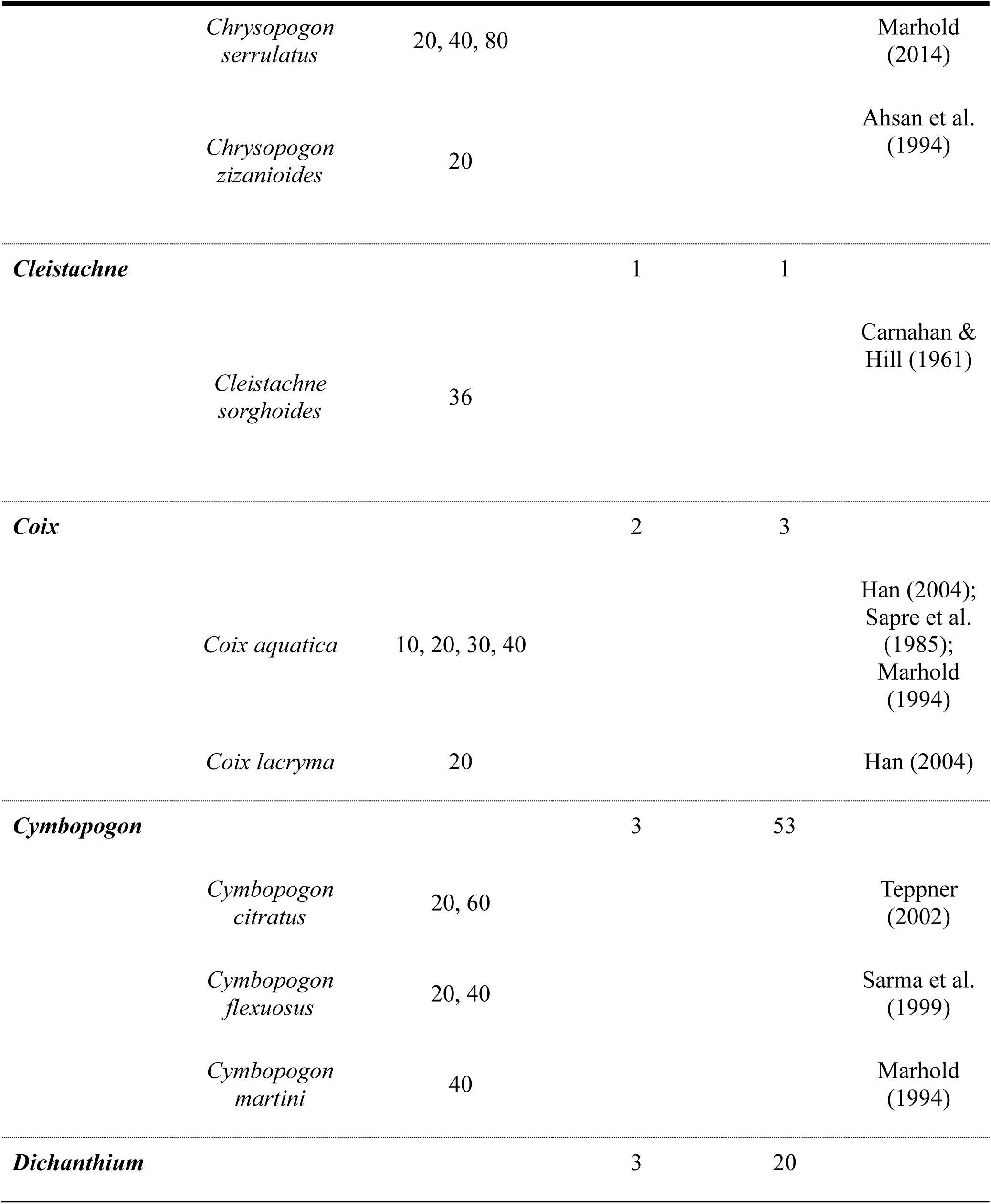
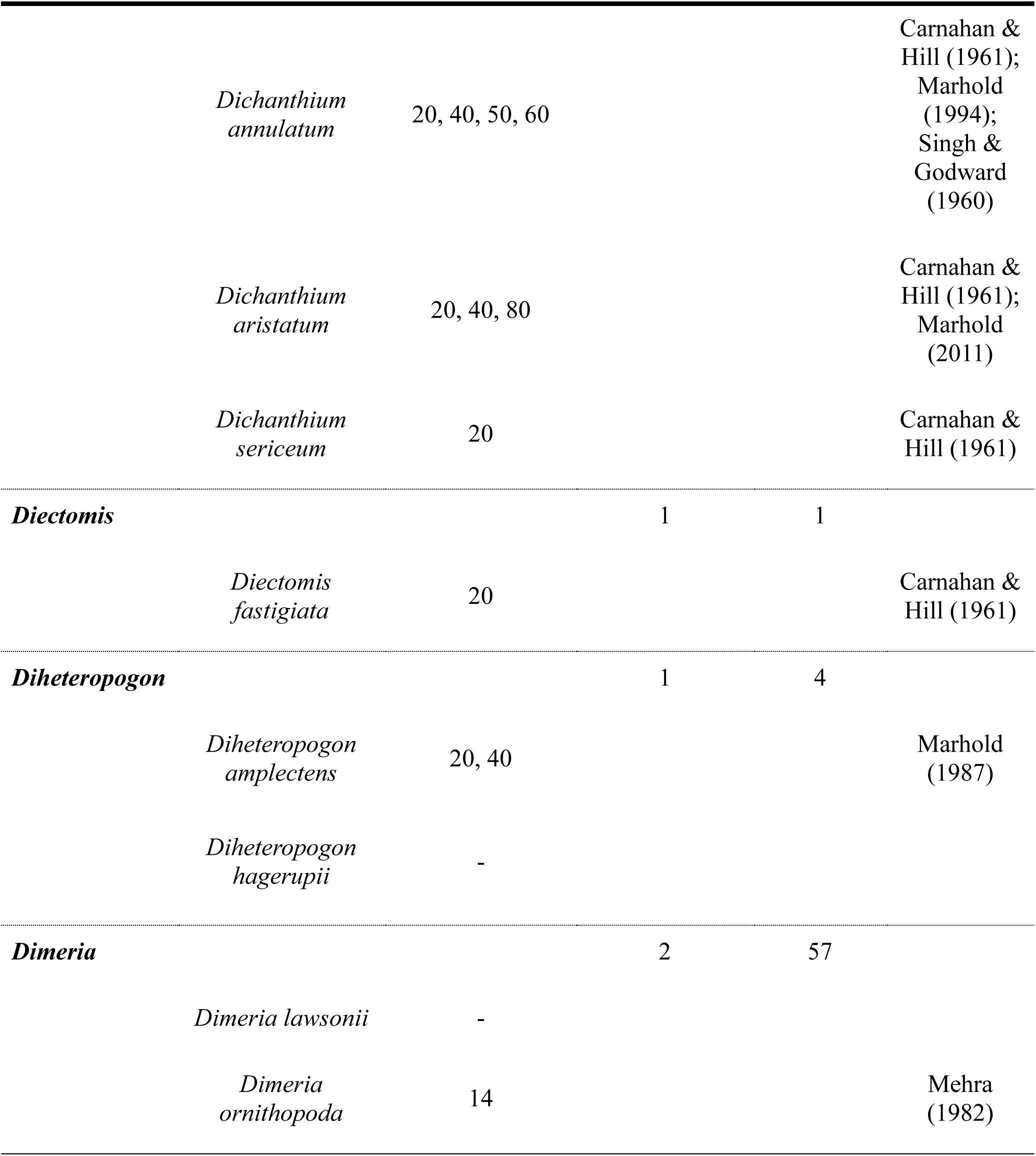
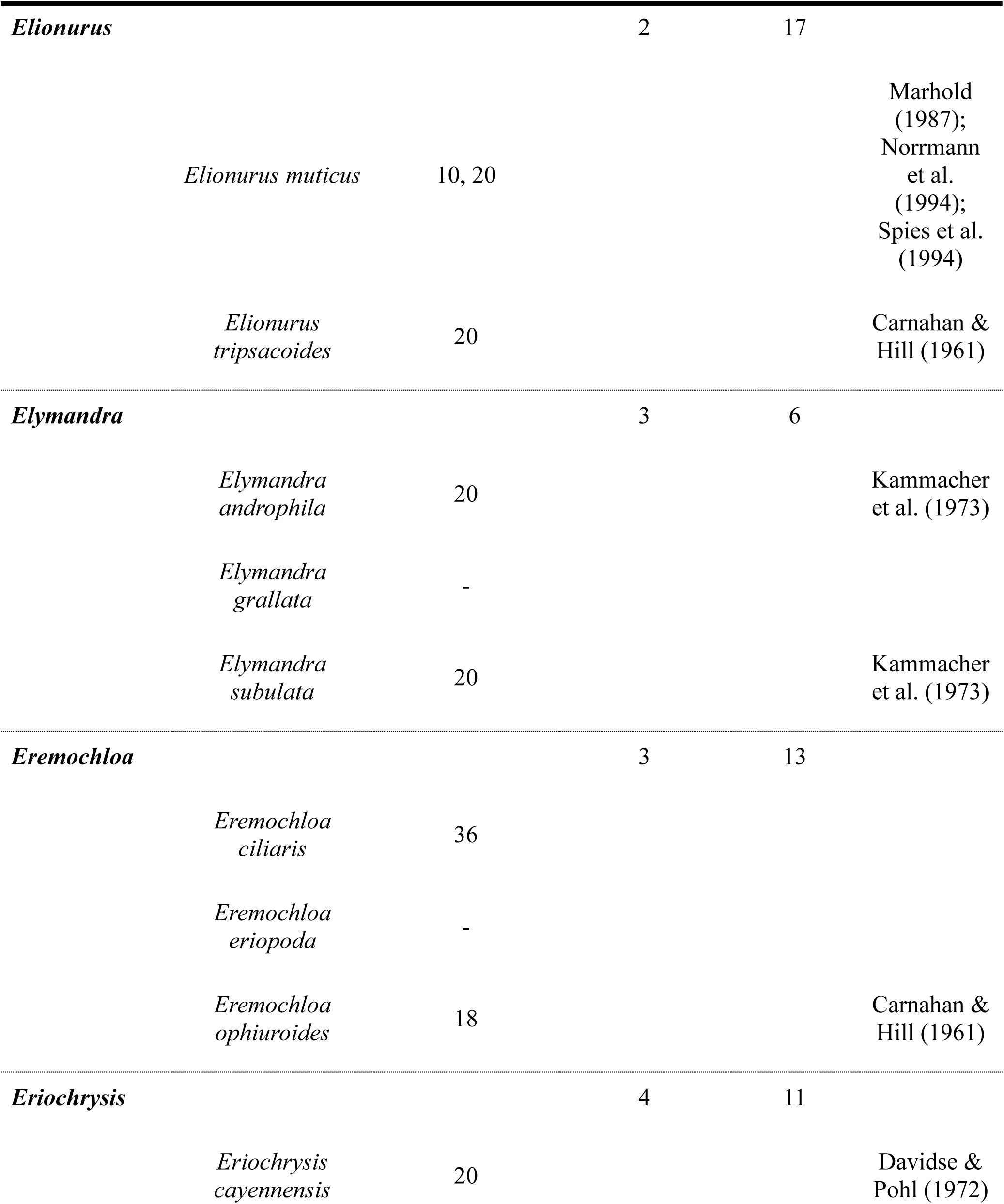
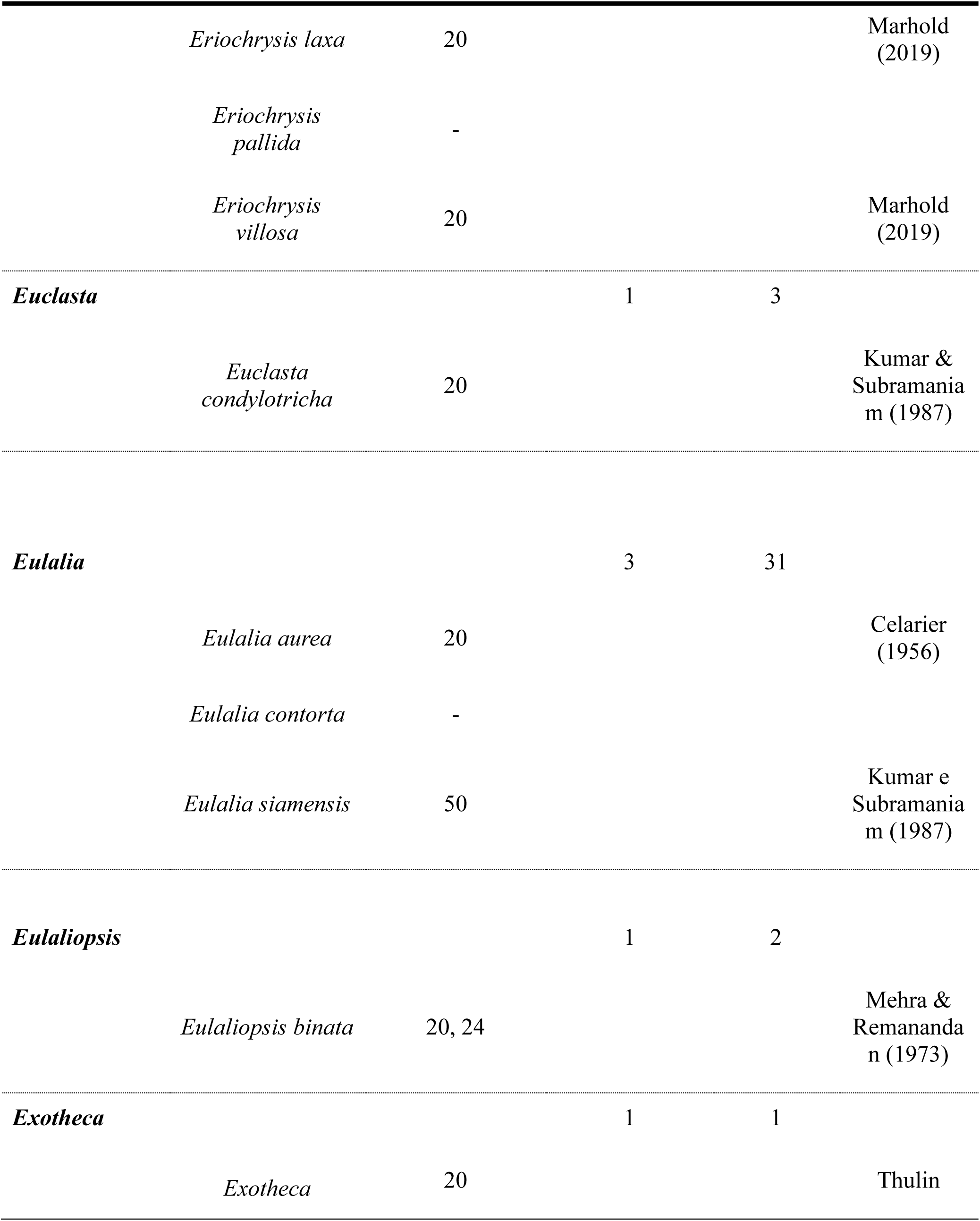
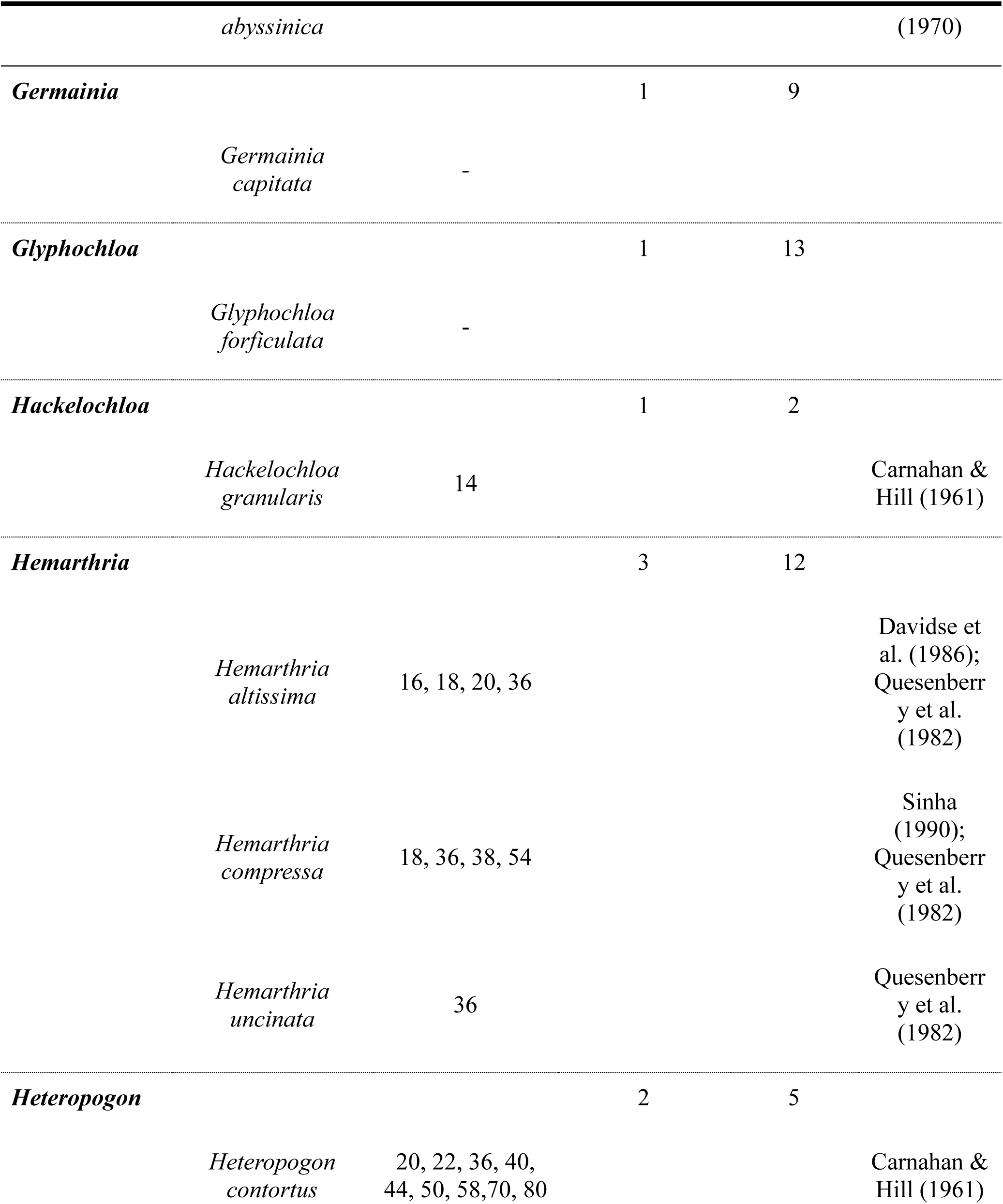
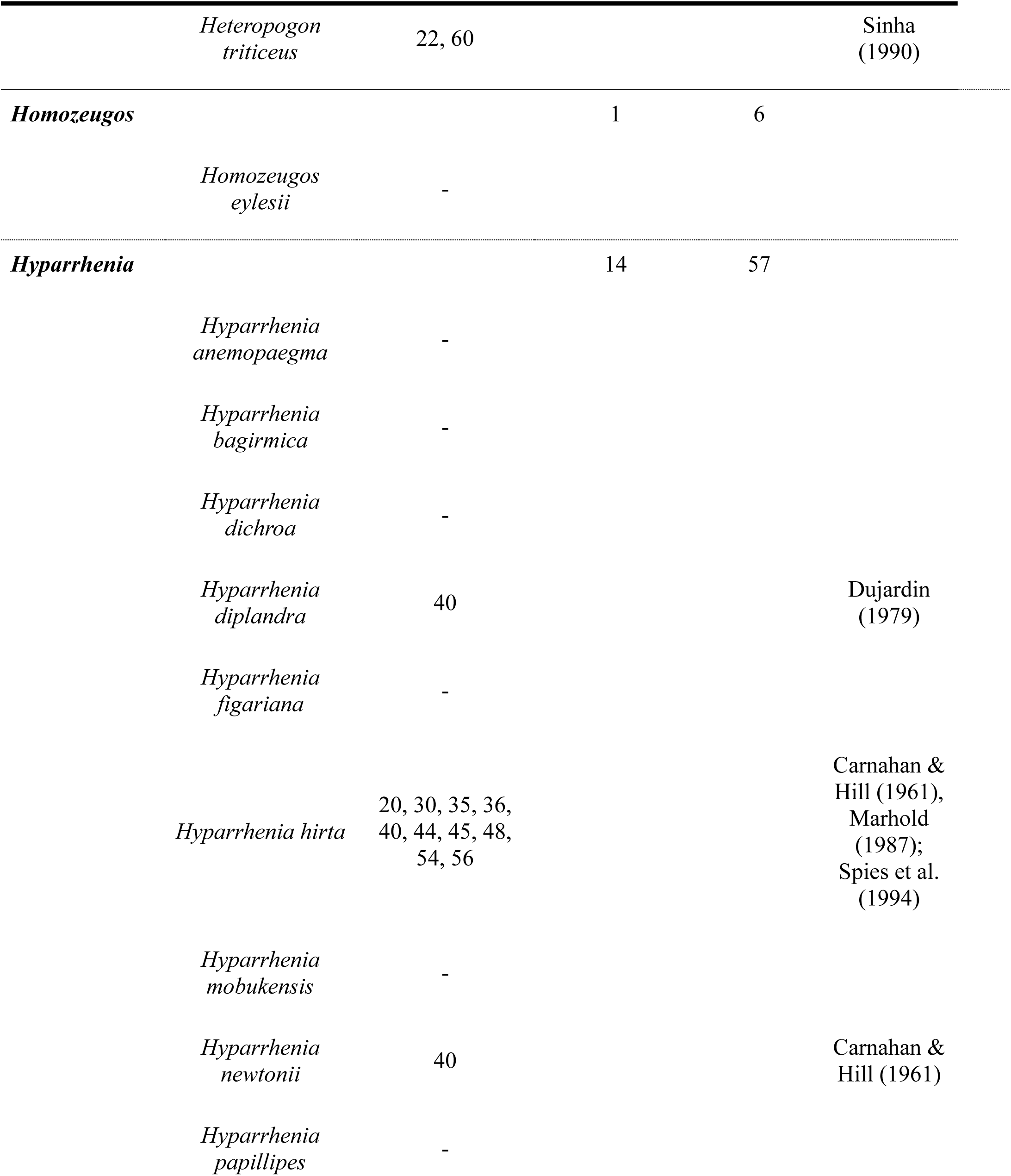
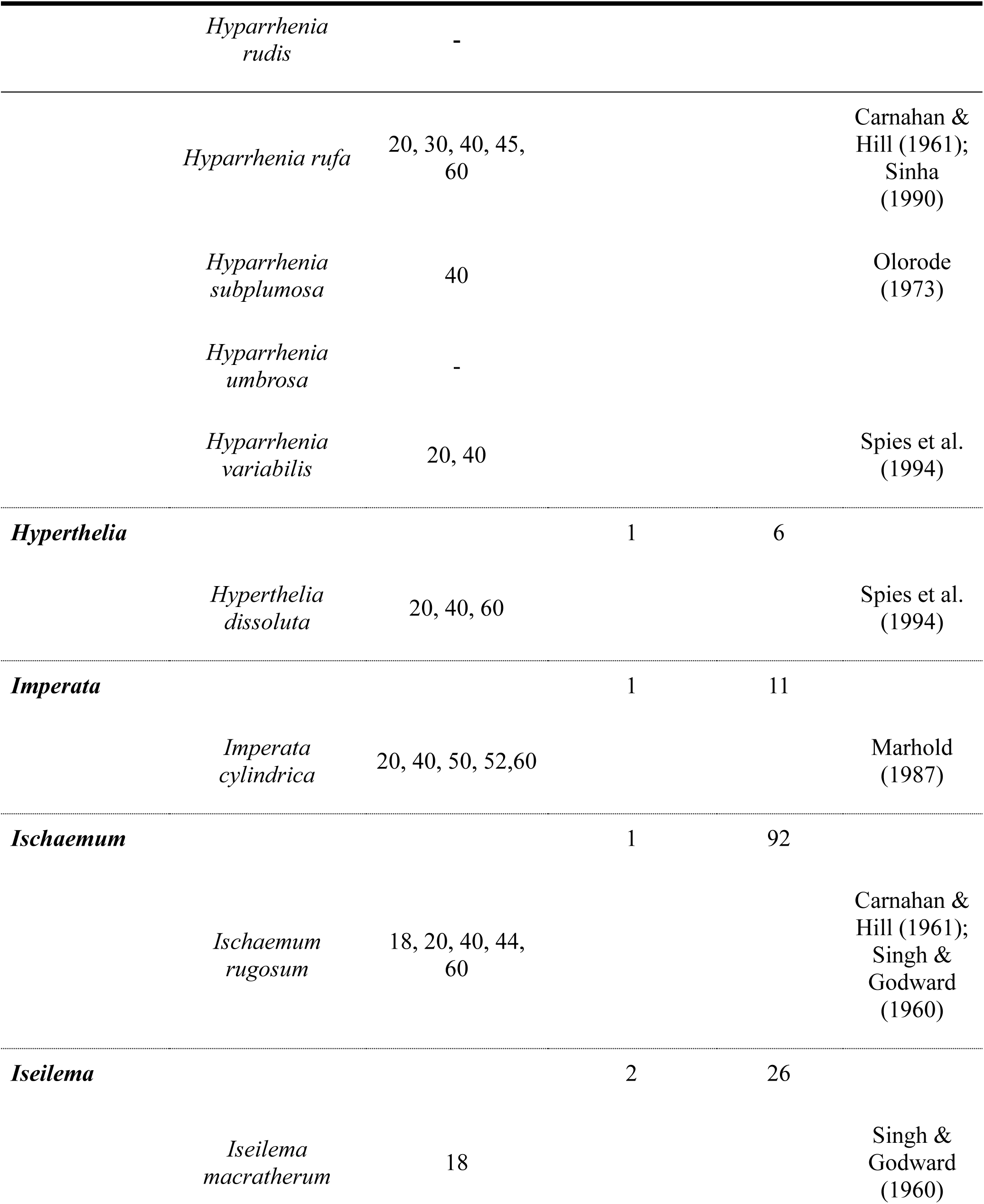
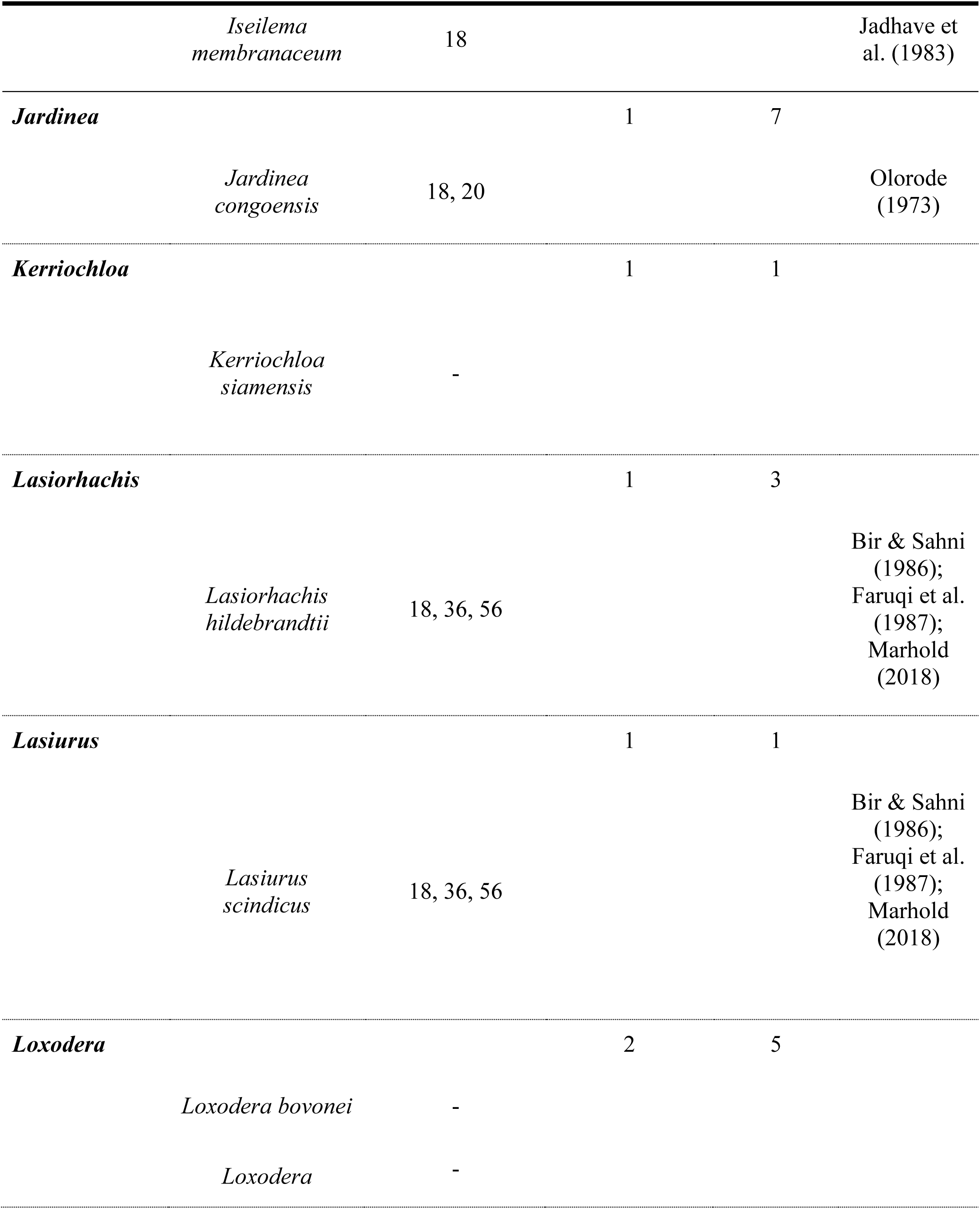
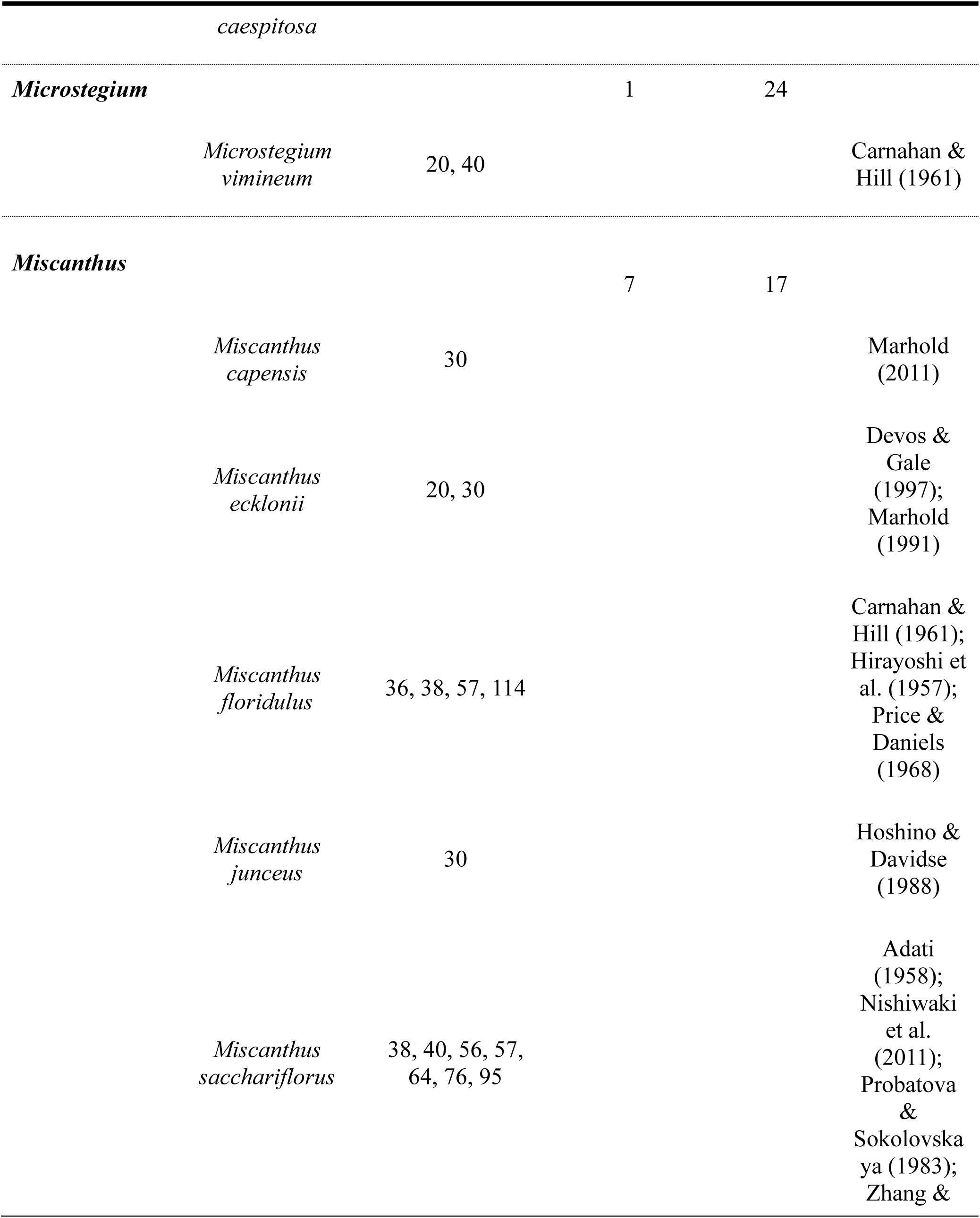
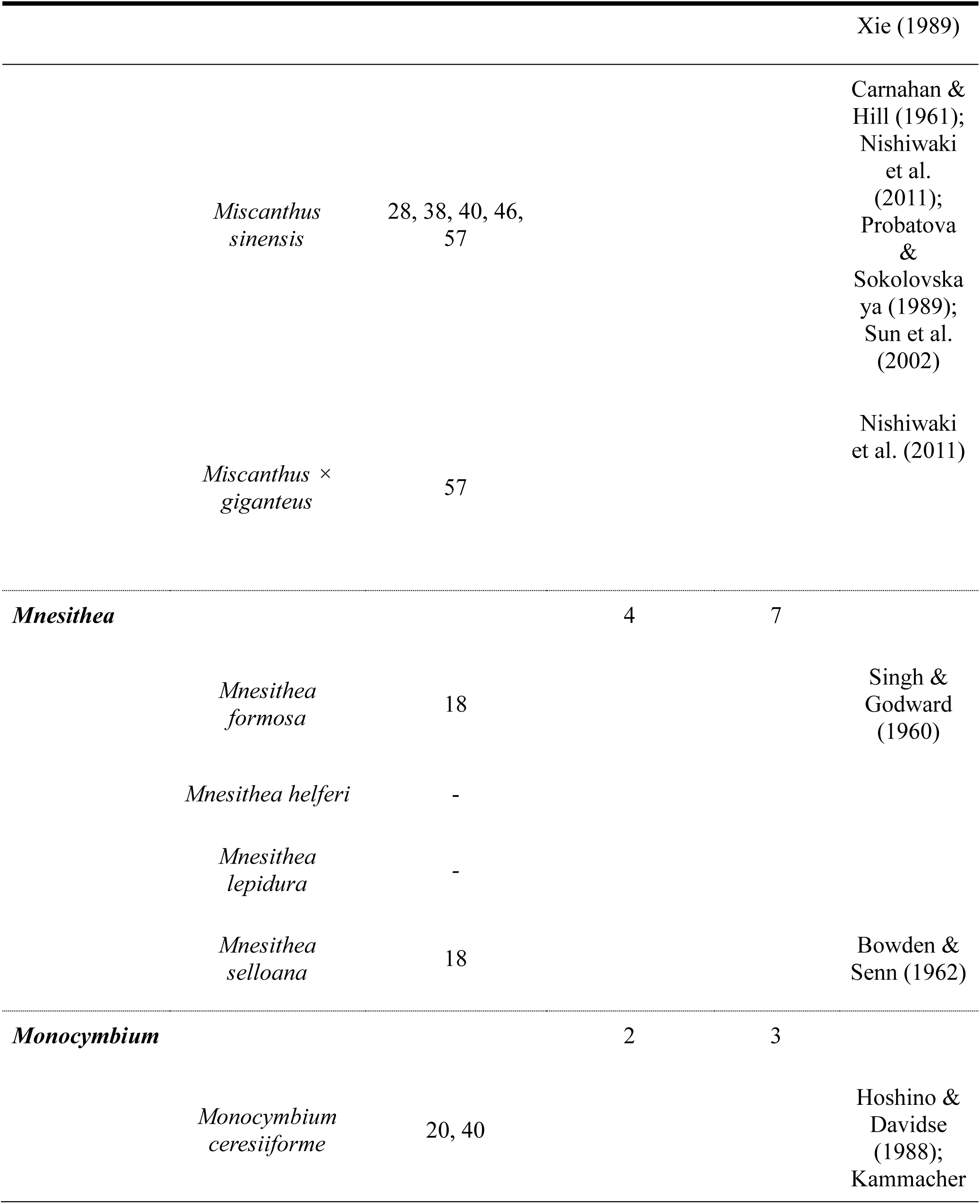
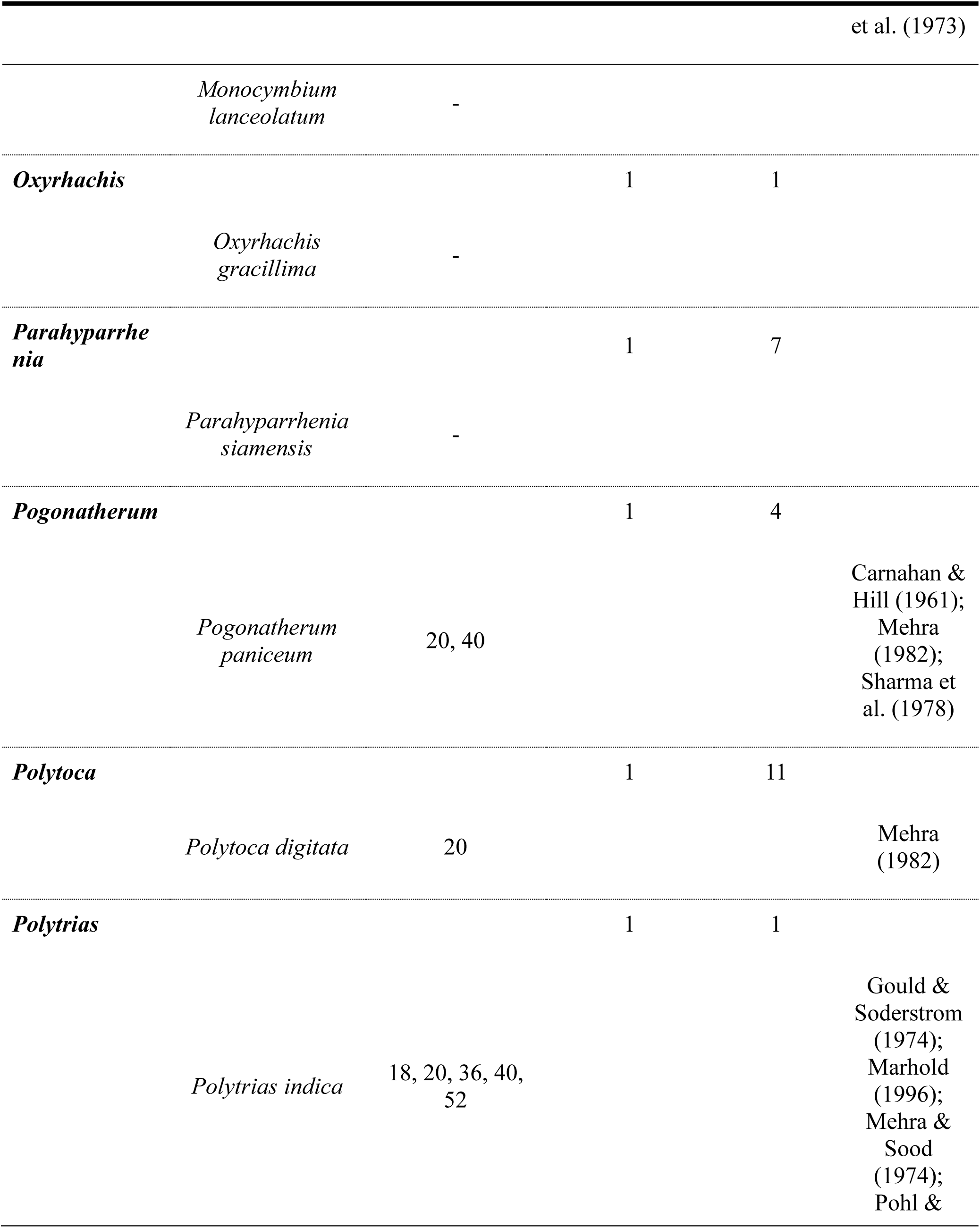
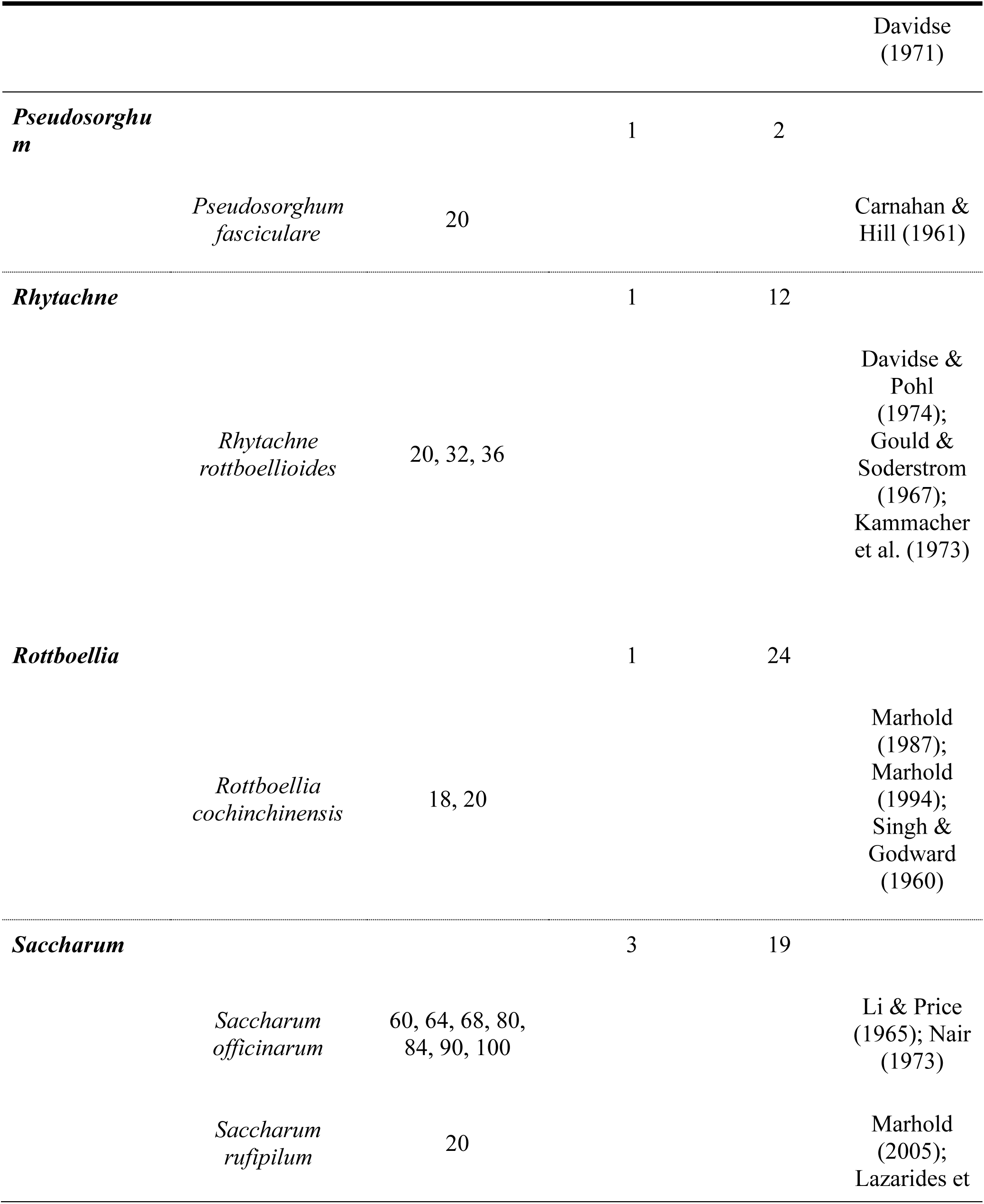
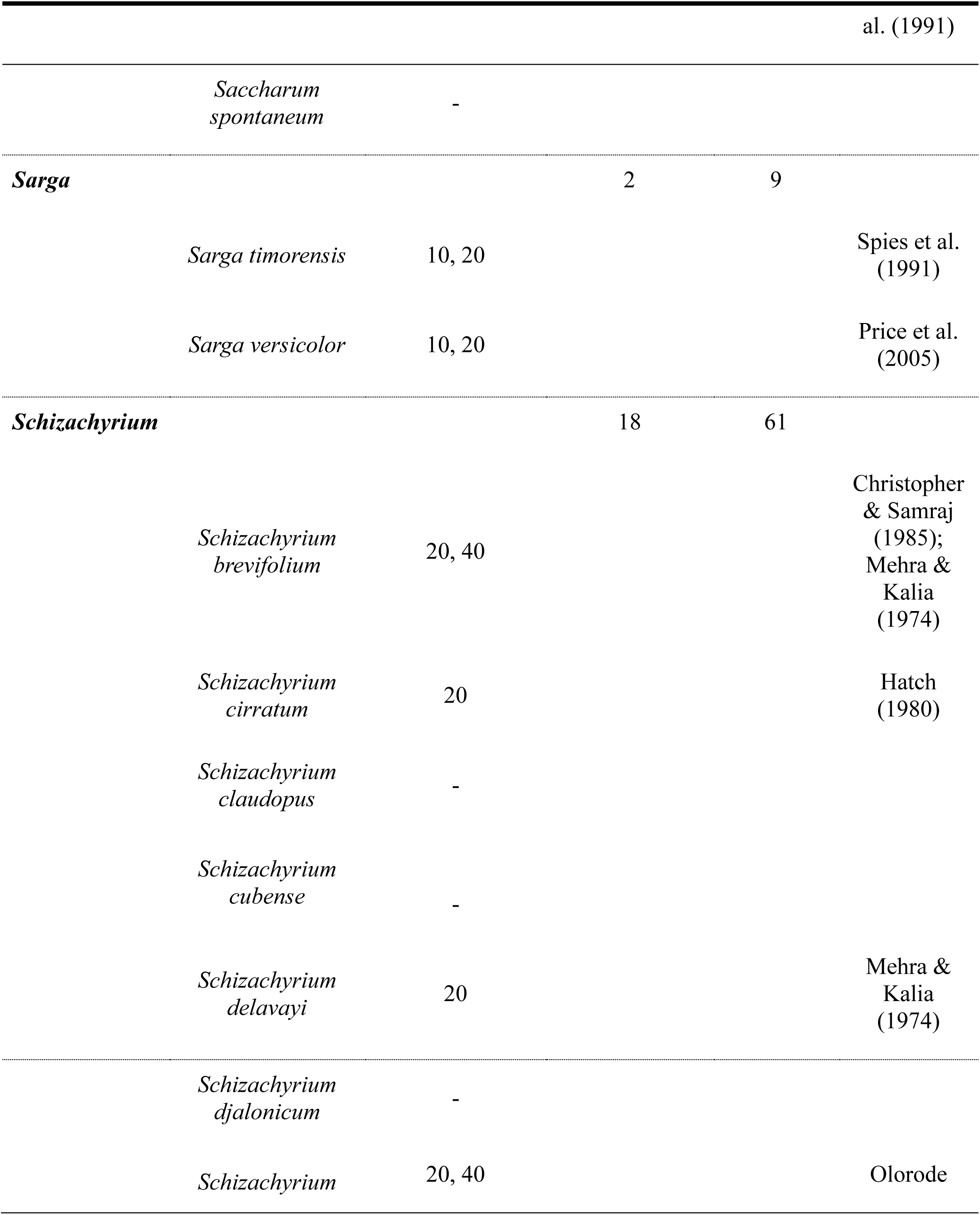
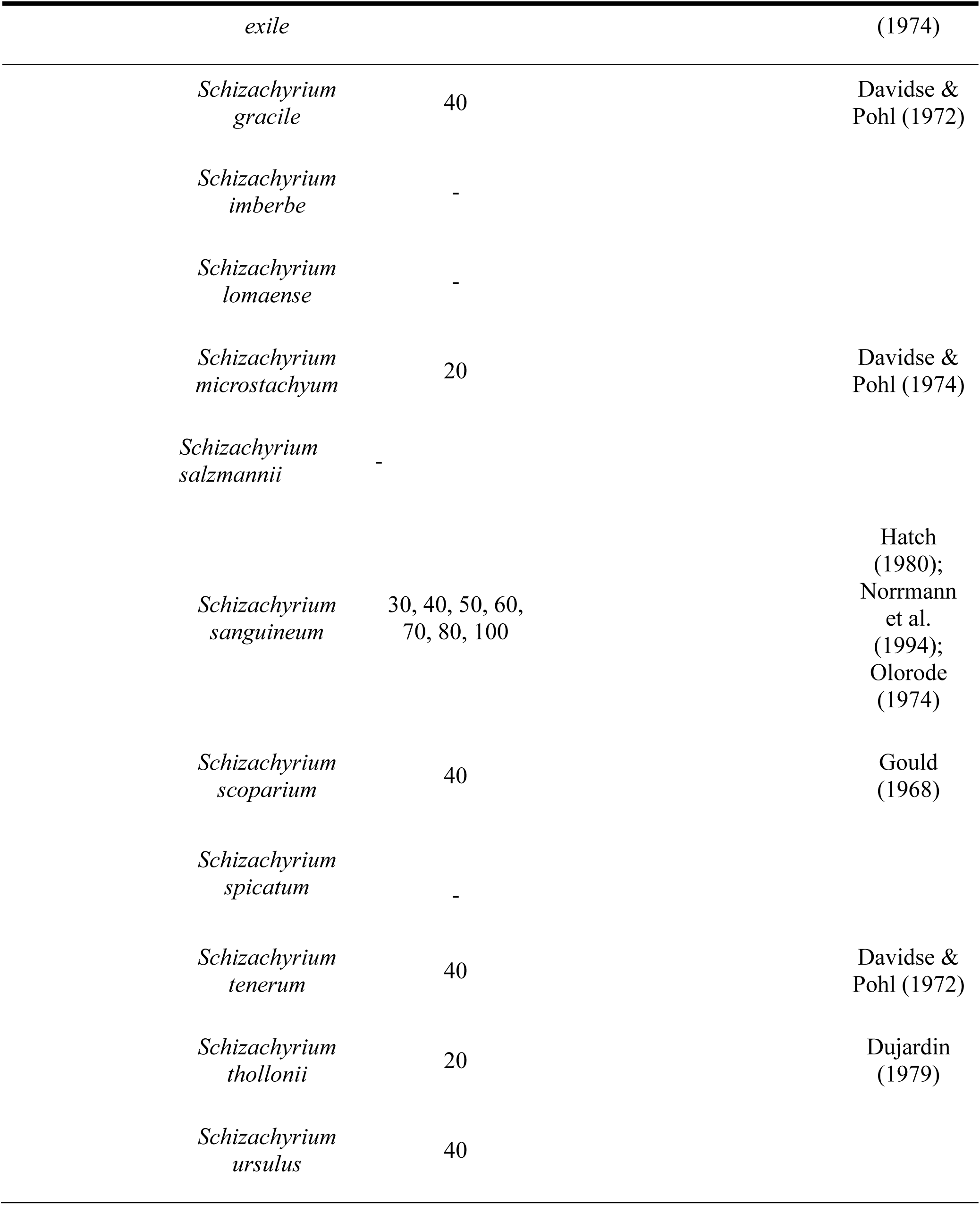
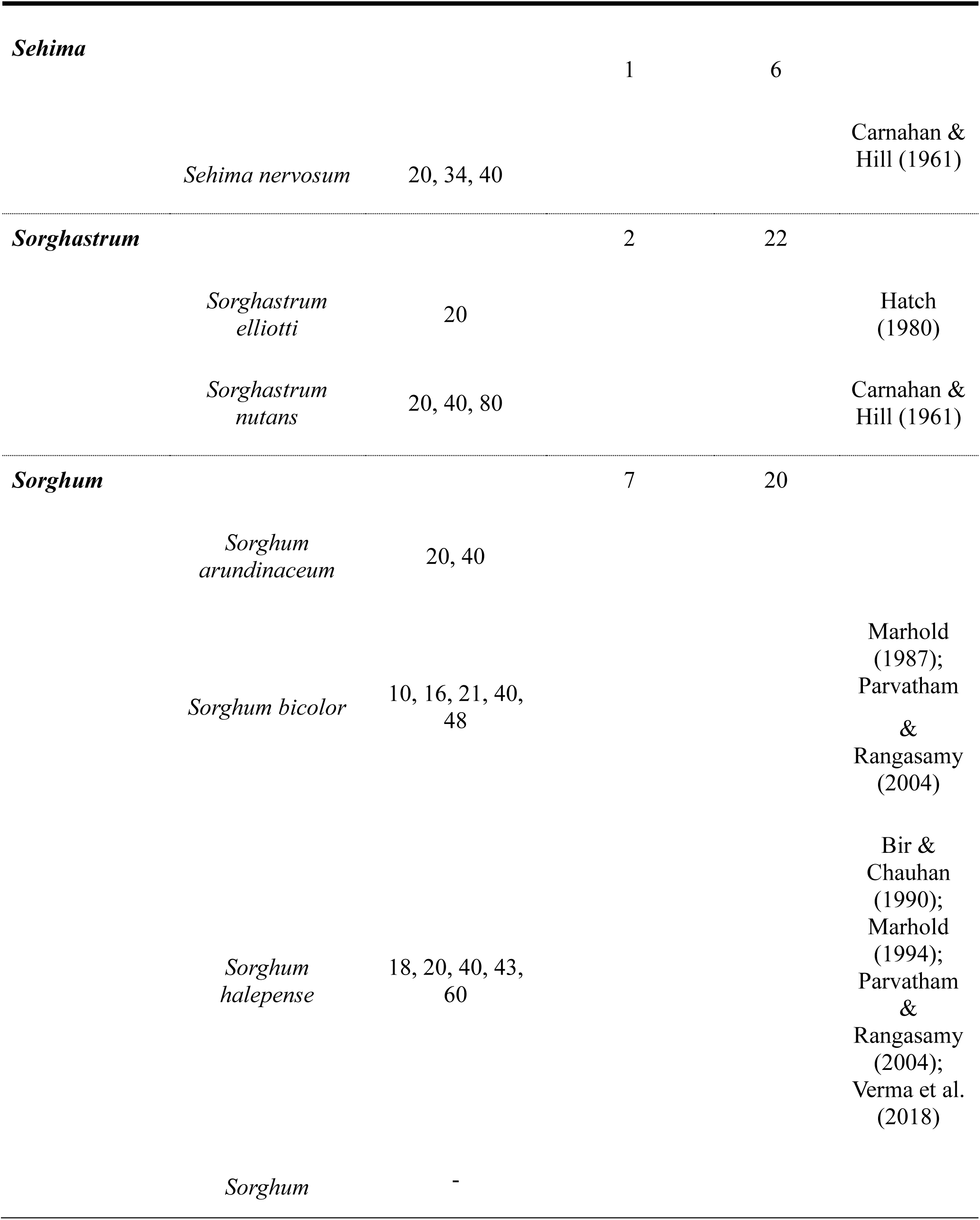
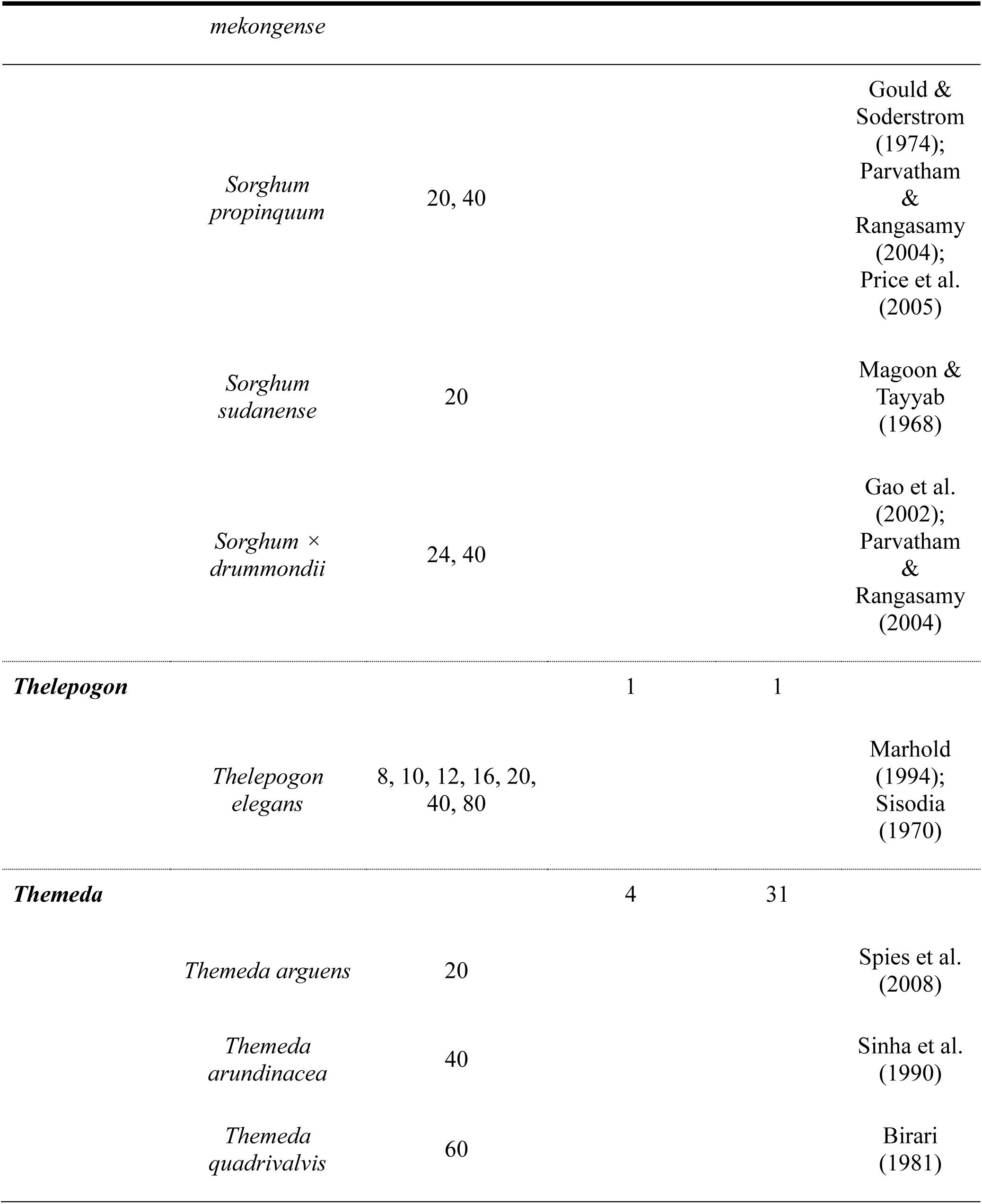
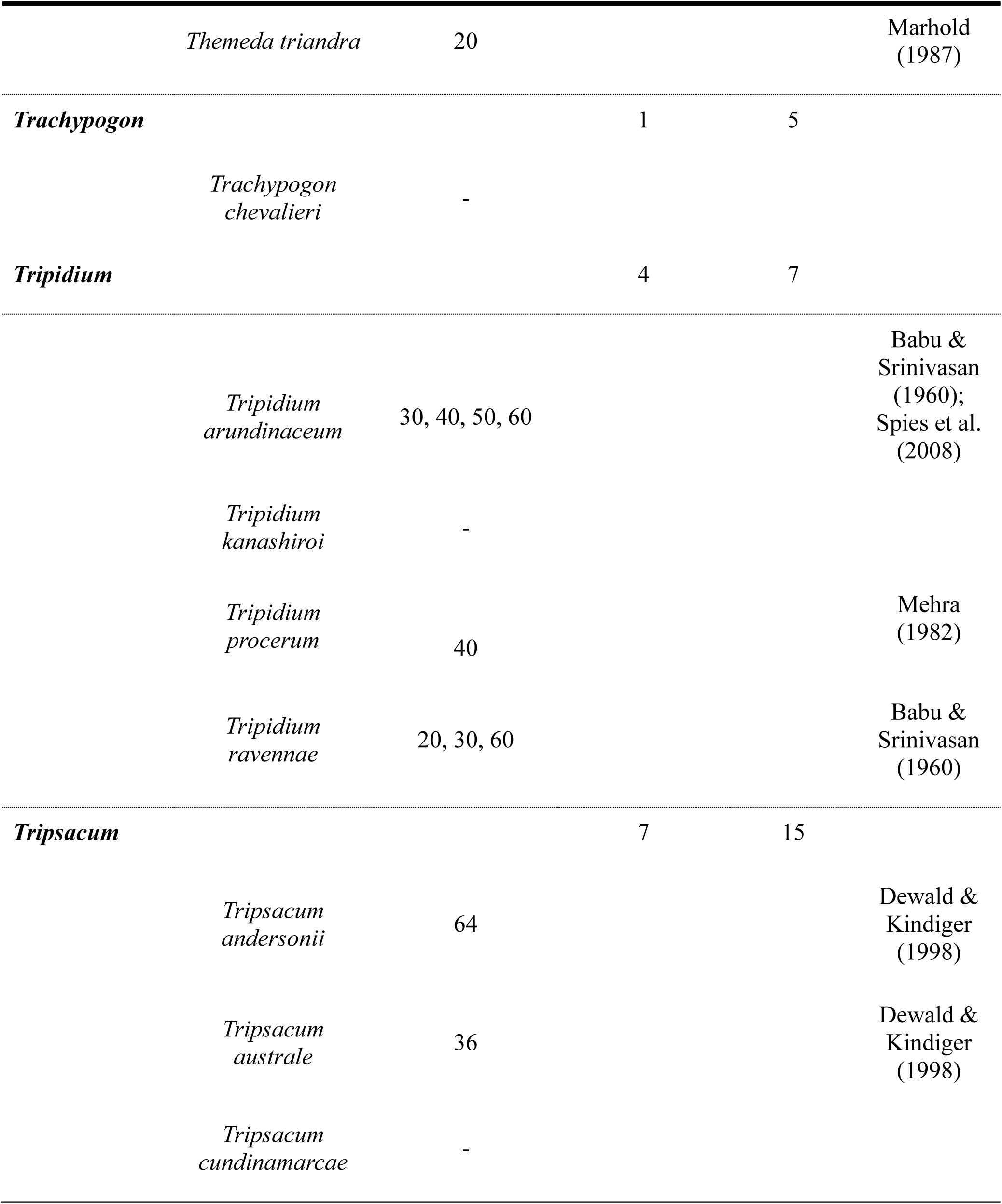
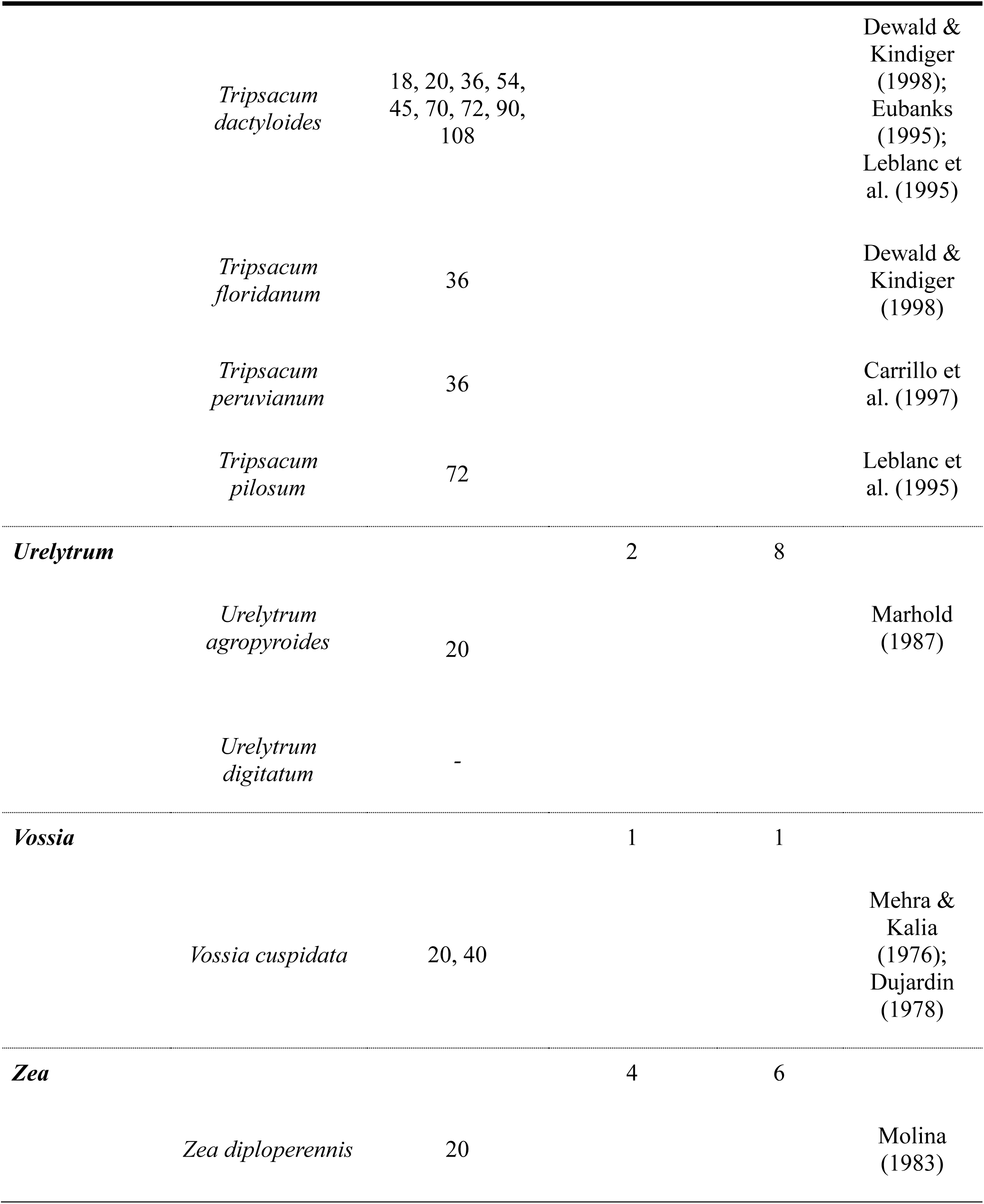
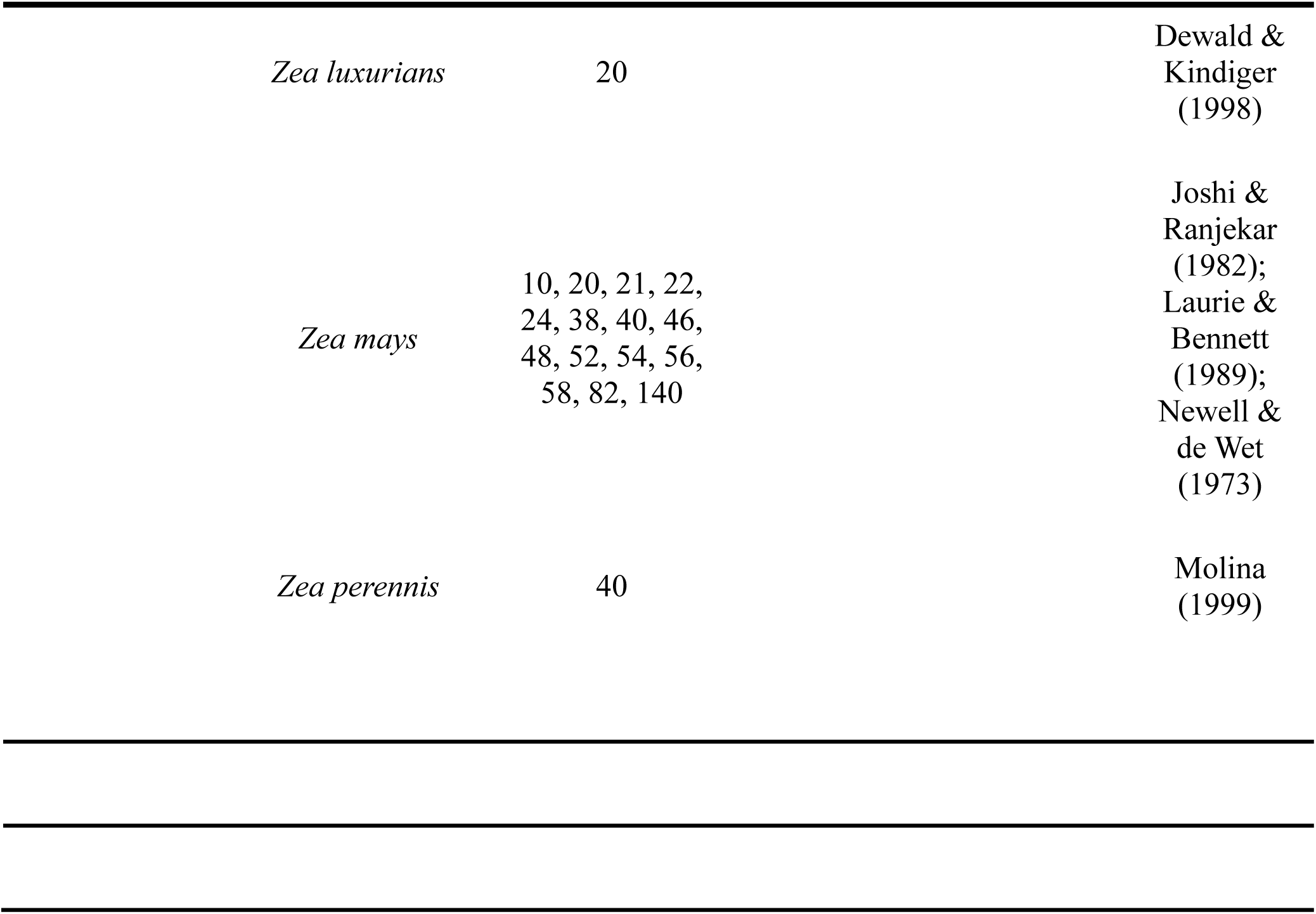
Chromosome numbers (2n) available in the literature for the selected species of the Andropogoneae tribe, according to the phylogenetic tree by Welker et al. (2020). The table also indicates the number of species from each genus included in this study, as well as the total number of species per genus, based on Welker et al. (2020) and the *Tropicos* platform.

**Table 2.**
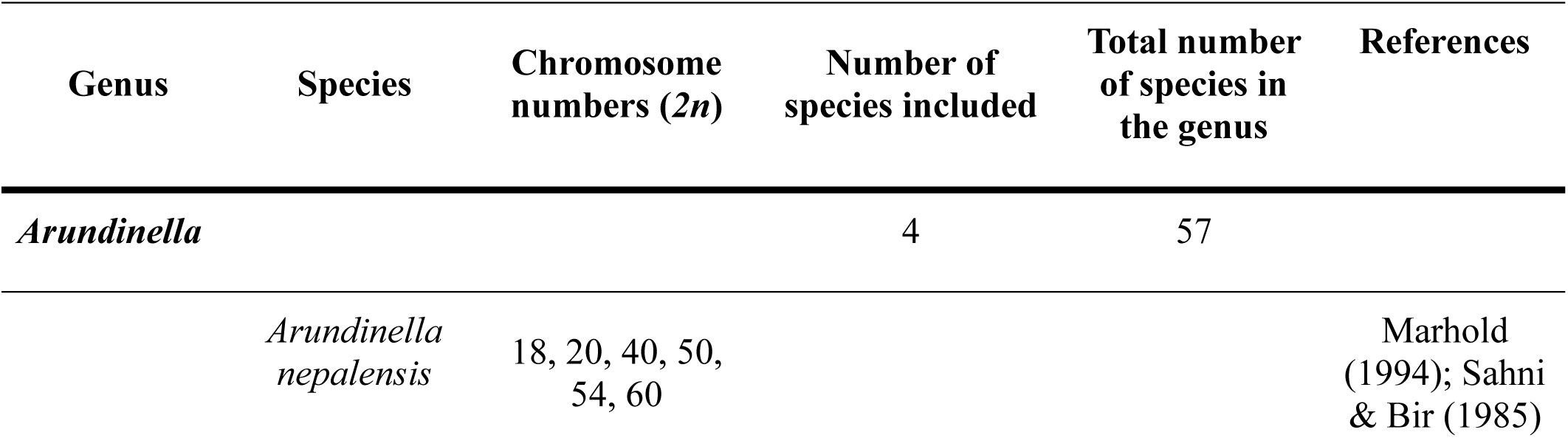
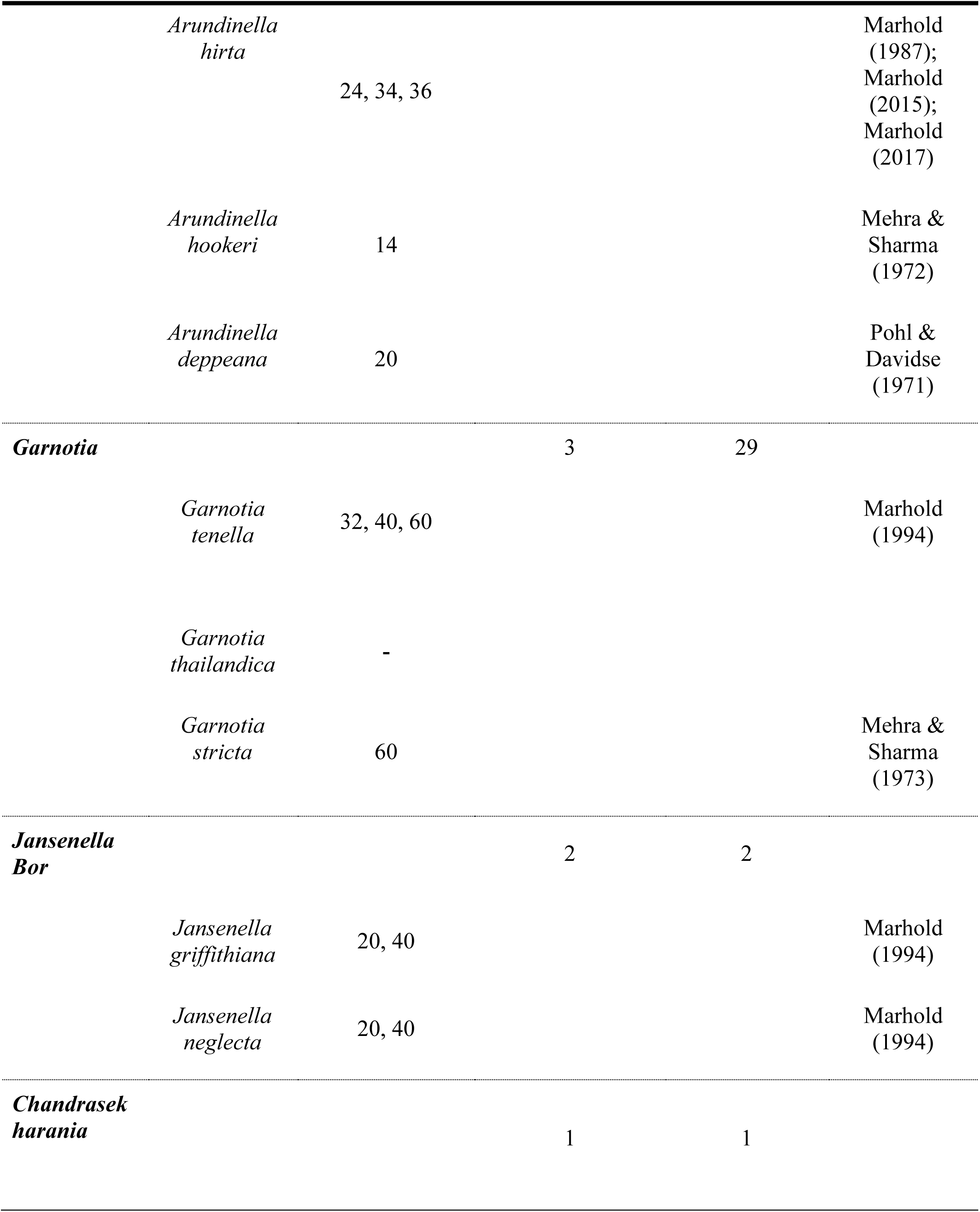
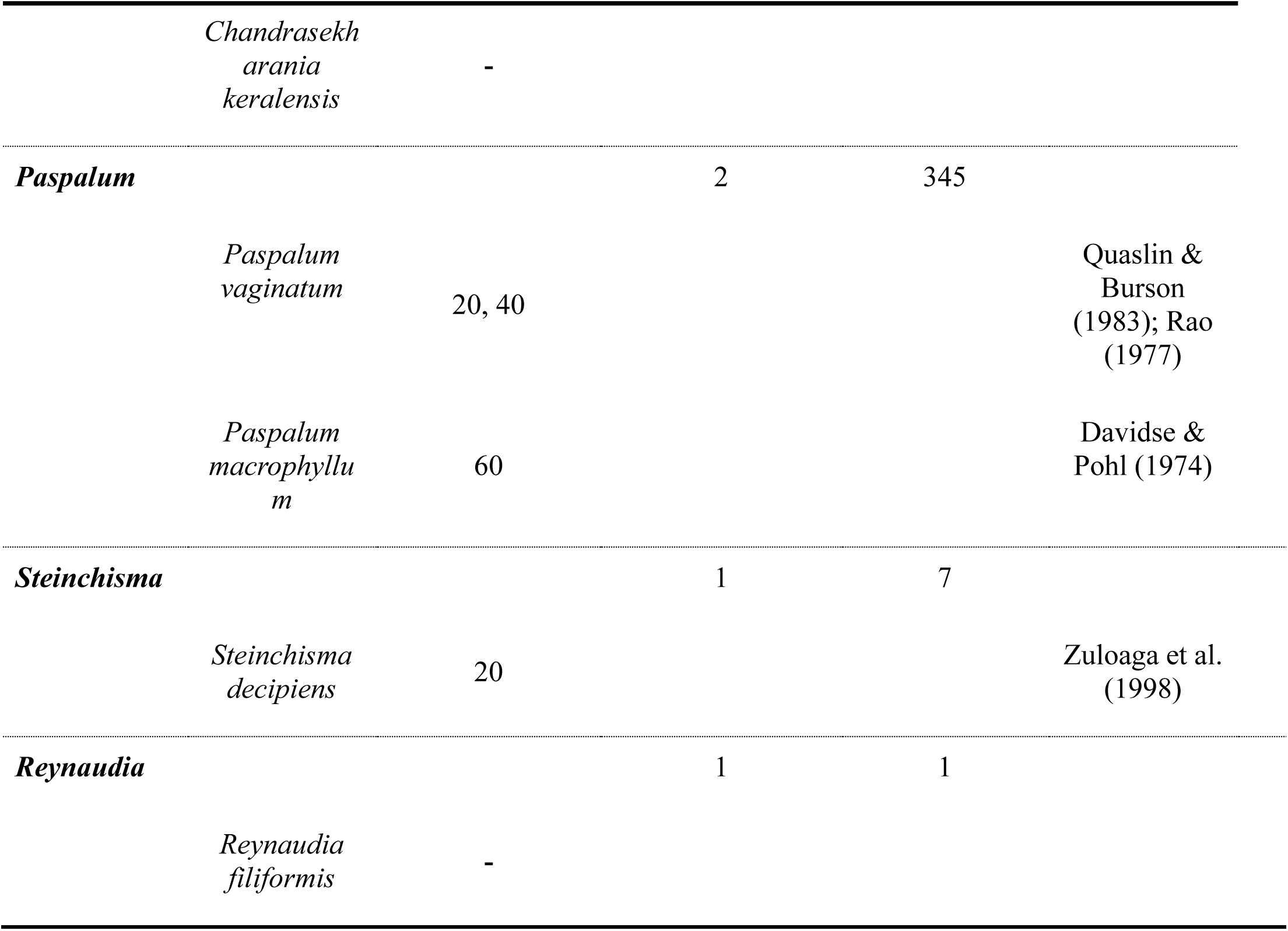
Chromosome numbers (2n) available in the literature for species of the Panicoideae subfamily that are phylogenetically close to the Andropogoneae tribe, according to the phylogenetic tree by Welker et al. (2020). The table also indicates the number of species from each genus included in this study, as well as the total number of species per genus, based on Welker et al. (2020).

The *2n* chromosome numbers found for the Andropogoneae tribe encompassed a wide range of values, from a minimum of *2n* = 8 (found in *Thelepogon elegans*) to a maximum of *2n* = 140 (observed in *Zea mays*). The value of *2n* = 20 was identified as the most frequent chromosome number in the data table. In the group of phylogenetically close species, the numbers ranged from *2n* = 14 (*Arundinella hookeri*) to *2n* = 60. The greatest variations in *2n* chromosome number within individual species were observed in *Zea mays* (with fifteen distinct chromosome numbers reported), *Hyparrhenia hirta* (ten), and *Bothriochloa bladhii*, *Heteropogon contortus*, and *Tripsacum dactyloides* (each with nine). Despite the extensive survey, chromosomal data were not found for 52 of the 230 listed species (about 23%), spanning 26 genera, with the largest gaps in the *Andropogon*, *Hyparrhenia*, and *Schizachyrium* genera.

### Data Analysis and Phylogenetic Tree

The Bayesian Inference utilized for the phylogenetic tree reconstruction (Figures 1a,b, c and d) confirmed robust evidence that the Andropogoneae tribe constitutes a monophyletic group.

**Fig. 1a.**
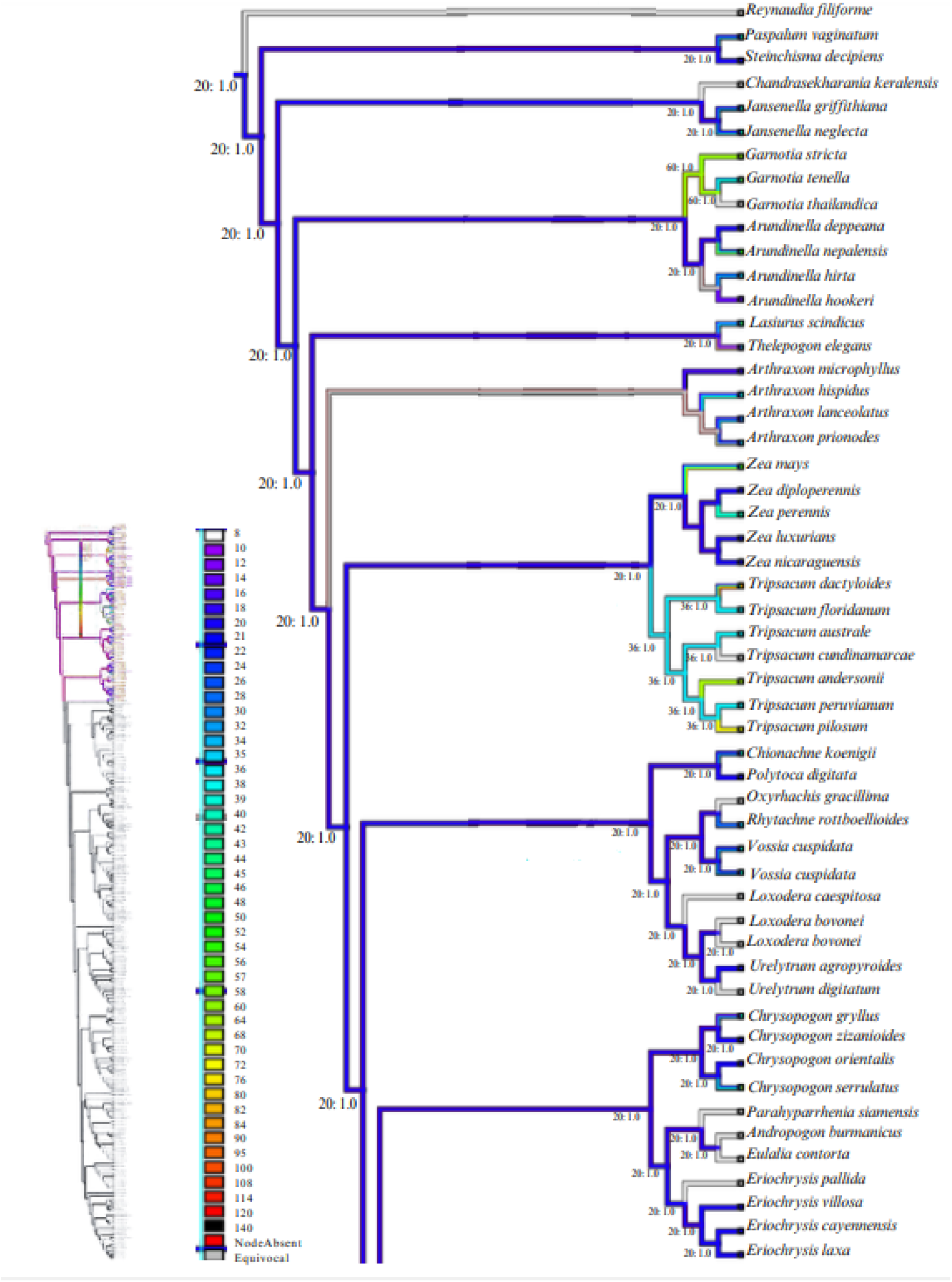
Phylogenetic tree of the Andropogoneae tribe with the reconstruction of chromosome number (*2n*) character states. The tree illustrates the evolutionary relationships among the species of the tribe and phylogenetically related species, according to Welker et al. (2020), highlighting the variation in chromosome numbers observed within each group.

**Fig. 1b.**
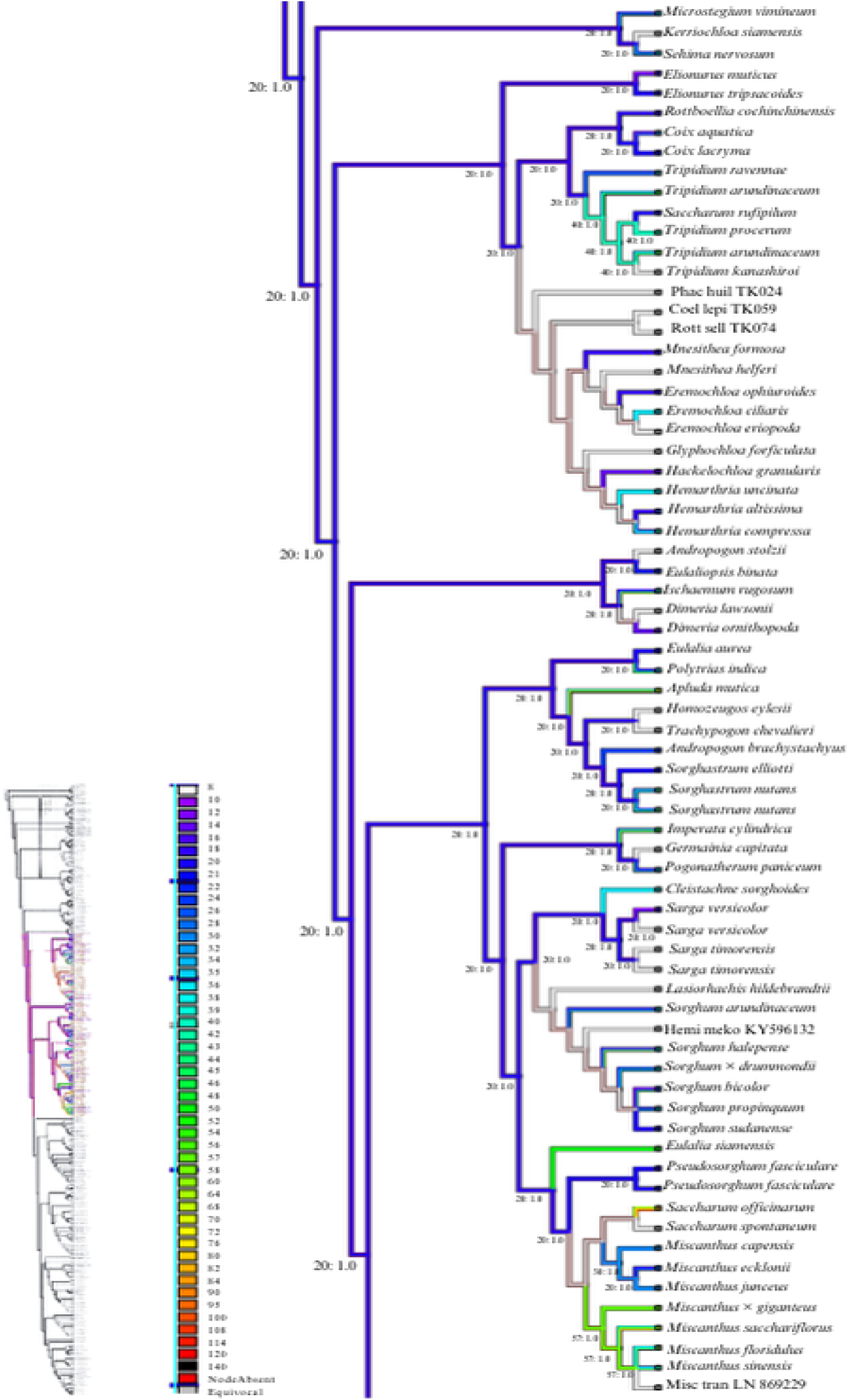
Phylogenetic tree of the Andropogoneae tribe with the reconstruction of chromosome number (*2n*) character states. The tree illustrates the evolutionary relationships among the species of the tribe and phylogenetically related species, according to Welker et al. (2020), highlighting the variation in chromosome numbers observed within each group.

**Fig. 1c.**
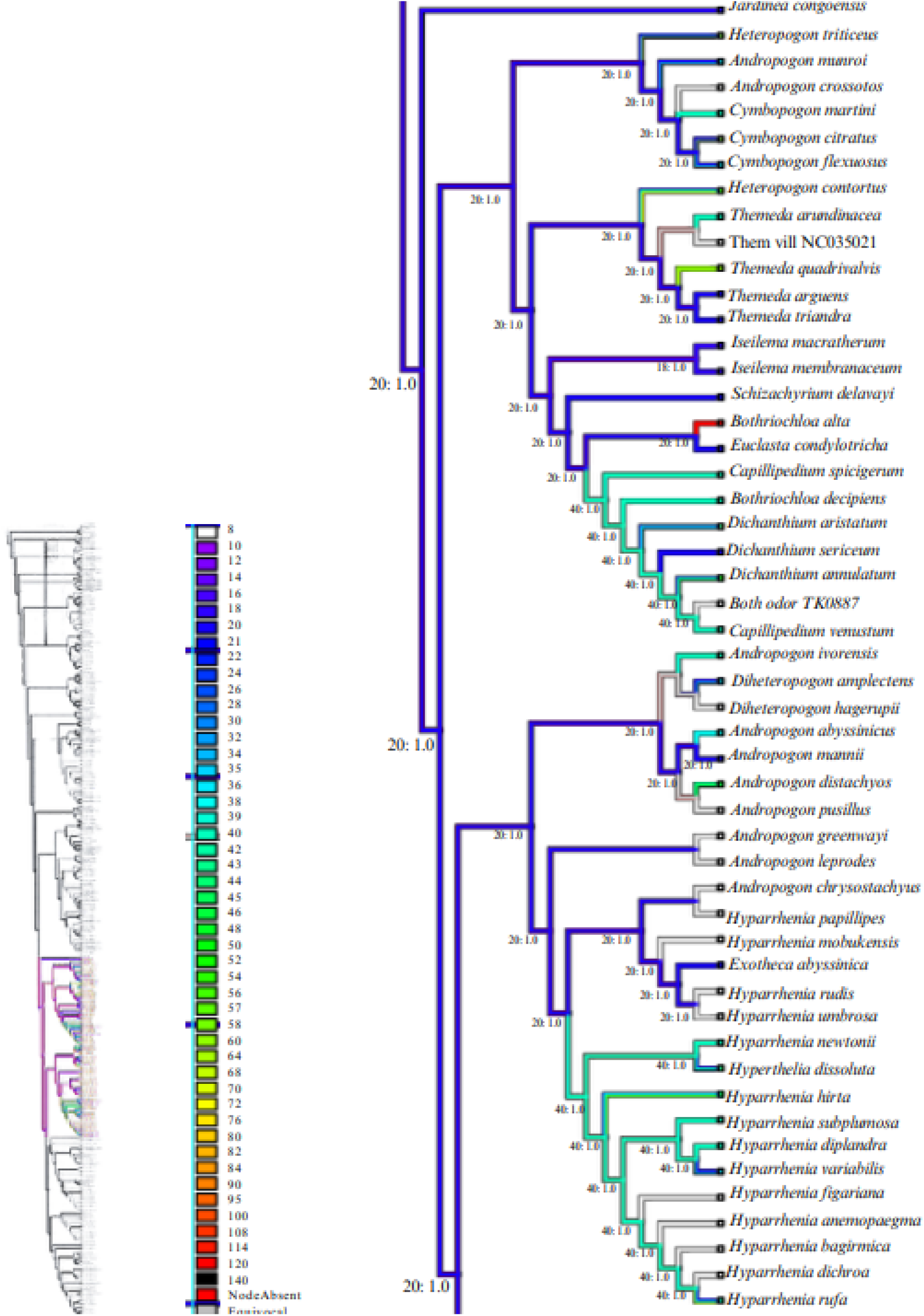
Phylogenetic tree of the Andropogoneae tribe with the reconstruction of chromosome number (*2n*) character states. The tree illustrates the evolutionary relationships among the species of the tribe and phylogenetically related species, according to Welker et al. (2020), highlighting the variation in chromosome numbers observed within each group.

**Fig. 1d.**
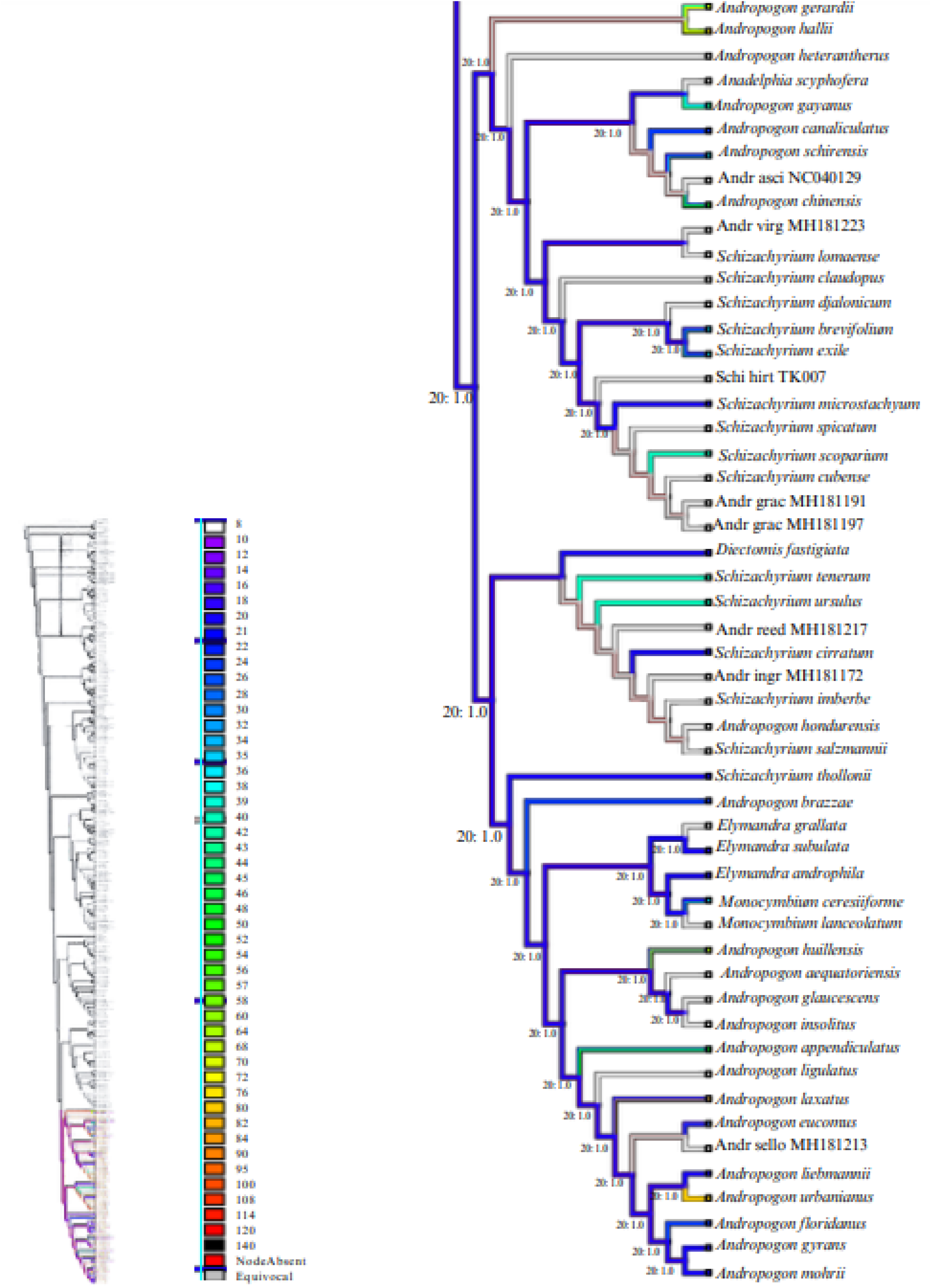
Phylogenetic tree of the Andropogoneae tribe with the reconstruction of chromosome number (*2n*) character states. The tree illustrates the evolutionary relationships among the species of the tribe and phylogenetically related species, according to Welker et al. (2020), highlighting the variation in chromosome numbers observed within each group.

The reconstruction of chromosomal characters revealed that the haploid number (*n* = 10) is the ancestral state for the majority of the internal nodes in the phylogeny, suggesting that this is the common primitive condition for many ancestral lineages. In contrast, the more terminal nodes of the tree, which represent more recently divergent taxa, consistently exhibit higher chromosome numbers. The variation in the *Zea* clade (from *2n* = 10 to *2n* = 140) exemplifies this trend, indicating that multiple polyploidization events and other chromosomal alterations occurred throughout the evolution of the more recent lineages, resulting in the wide range of observed numbers.

Specifically for the Andropogoninae subtribe (which includes genera such as *Andropogon*, *Hyparrhenia*, and *Schizachyrium*), the phylogenetic analysis demonstrates a consistent predominance of the *2n* = 20 chromosome number across its branches. This predominance suggests that *2n* = 20 may be the ancestral or primitive state for most genera within Andropogoninae. The notable stability and cytogenetic conservation of this number, despite the diversity and adaptation observed in other groups, suggests a strong link between this genomic structure and the subtribe’s adaptive success, pointing to a common ancestor that preserved this essential characteristic.

## Discussion

### Ancestral chromosome number and evolutionary stability

The phylogenetic reconstruction confirms that *2n* = 20 (x = 10) represents the ancestral chromosome number for tribe Andropogoneae, consistent with previous hypotheses based on limited taxonomic sampling (Spangler et al., 1999; Giussani et al., 2001; Price et al., 2005). This ancestral state has been maintained across most major lineages, indicating strong stabilizing selection or developmental constraints favoring this chromosome number (Stebbins, 1971). The predominance of *2n* = 20 at internal nodes throughout the phylogeny suggests that this karyotype configuration provides optimal genomic organization for Andropogoneae species (Clayton & Renvoize, 1986).

The basic chromosome number x = 10 appears to be plesiomorphic not only for Andropogoneae but potentially for much of Panicoideae, as evidenced by its occurrence in phylogenetically related tribes including Arundinelleae and Paspaleae (Soreng et al., 2022; Welker et al., 2020). Spangler et al. (1999) and Wilson et al. (1999) suggested that x = 10, rather than x = 5, represents the base number of the tribe based on the wide variety of taxa near the base of their phylogenetic trees exhibiting *n* = 10. This chromosomal conservation across deep evolutionary timescales contrasts sharply with the extensive variation observed in more recently diverged lineages, highlighting the dynamic nature of chromosomal evolution in grasses (Kellogg, 2015; Elliott et al., 2022).

### Polyploidy as a driver of diversification

Approximately 30% of Andropogoneae species examined in this study exhibited polyploidy, consistent with previous estimates that one-third of tribe species originated through polyploidization events (Estep et al., 2014). Polyploidy, particularly allopolyploidy arising from interspecific hybridization, has been a major driver of speciation and diversification within Andropogoneae (Guerra, 2008; Peer et al., 2020). This finding aligns with broader patterns across vascular plants, where approximately 35% of species have undergone whole-genome duplication (Peer et al., 2020; Leitch & Leitch, 2013).

Multiple polyploid series were evident across numerous genera. Genus *Andropogon* exemplifies this pattern, with species exhibiting *2n* = 20 (diploid), *2n* = 40 (tetraploid), *2n* = 60 (hexaploid), *2n* = 80 (octoploid), and even *2n* = 140 (14-ploid) (Carnahan & Hill, 1961; Campbell, 1983; Nagahama & Norrmann, 2012). Such extensive polyploid series suggest recurrent polyploidization events, potentially involving both autopolyploidy and allopolyploidy (Spies et al., 1994; Wood, 2009). The presence of intermediate chromosome numbers (*2n* = 68, 70 in *A. gerardii*) may reflect aneuploidy following polyploidization, a common phenomenon in polyploid complexes (Mayrose et al., 2010; Schubert & Lysak, 2011).

Economically important genera showed particularly high polyploidy frequencies. In genus Saccharum, *S. officinarum* (cultivated sugarcane) exhibited *2n* = 60, 64, 68, 80, 84, 90, and 100, reflecting complex allopolyploid origins involving multiple progenitor species (Daniels & Roach, 1987). Similarly, genus Miscanthus, increasingly important for bioenergy production, displayed extensive chromosomal variation with multiple polyploid levels (Hodkinson et al., 2002). The triploid hybrid M. × giganteus (*2n* = 57) exemplifies allopolyploidy’s role in crop development, combining favorable traits from diploid and tetraploid parents.

Genus *Zea* presents a particularly interesting case. While wild relatives generally exhibit *2n* = 20 or 40, cultivated maize (Z. mays) displays remarkable chromosomal variation including *2n* = 10, 20, 40, and higher numbers up to 140 (Joshi & Ranjekar, 1982). The occurrence of *n* = 10 in some maize populations suggests descending dysploidy from the ancestral x = 10 diploid state, while higher numbers reflect polyploidization and aneuploidy associated with domestication and breeding.

### Phylogenetic patterns and biogeographic implications

Chromosomal variation in Andropogoneae exhibits clear phylogenetic structure, with related species often sharing similar chromosome numbers or belonging to polyploid series derived from common ancestors (Welker et al., 2020). Within subtribe Andropogoninae, the predominance of *2n* = 20 across genera including *Andropogon*, *Schizachyrium*, *Hyparrhenia*, and *Elymandra* indicates retention of the ancestral state in this diverse lineage (Sanchez-Ken & Clark, 2010; Kellogg, 2015).

However, certain clades show elevated rates of chromosomal evolution. The subtribe Tripsacinae, comprising genera *Tripsacum* and *Zea*, exhibits extensive chromosomal variation potentially associated with adaptation to temperate environments and agricultural development. Similarly, subtribe Saccharinae (*Miscanthus*, *Saccharum*) displays high polyploidy frequencies that may be related to their predominantly Asian distribution and adaptation to diverse ecological niches (Hodkinson et al., 2002).

The pantropical distribution of Andropogoneae and its ecological dominance in grassland ecosystems may be partly attributable to chromosomal variation facilitating ecological diversification (Sanchez-Ken & Clark, 2010; Estep et al., 2014). Polyploidy confers advantages including increased genetic variation, heterosis, and enhanced tolerance to environmental stress (Levin, 1993; Peer et al., 2020). These attributes may have enabled polyploid Andropogoneae lineages to colonize diverse habitats and respond to climatic fluctuations throughout their evolutionary history (Wood, 2009; Elliott et al., 2022).

### Knowledge gaps and future directions

Despite comprehensive literature survey, chromosomal data remain unavailable for numerous Andropogoneae species, limiting resolution of evolutionary patterns. Approximately 40% of sampled species lacked published chromosome counts, with particularly poor representation in tropical African and Asian taxa. This geographic bias in cytogenetic sampling impedes understanding of chromosomal evolution across the tribe’s entire distribution.

Future research should prioritize cytogenetic characterization of undersampled lineages, particularly within diverse genera such as *Andropogon* (135 species), *Ischaemum* (92 species), and *Cymbopogon* (53 species), for which chromosome numbers are known for fewer than 30% of species (Soreng et al., 2022). Integration of chromosome counts with genome size estimates, karyotype analysis, and molecular cytogenetic techniques such as fluorescent *in situ* hybridization (FISH) would provide deeper insights into genome evolution (Guerra, 2008).

The role of allopolyploidy versus autopolyploidy in Andropogoneae diversification remains poorly understood (Estep et al., 2014; Kim et al., 2014). Molecular phylogenetic analyses integrating nuclear gene sequences can distinguish these polyploid origins and identify parental species involved in allopolyploid formation. Such studies would clarify mechanisms underlying polyploid speciation and illuminate hybrid origins of economically important species including sugarcane and certain bioenergy grasses (Hodkinson et al., 2002).

The present findings emphasize the dynamic nature of chromosomal evolution in Andropogoneae, with polyploidy and dysploidy generating extensive variation upon which natural and artificial selection act (Schifino-Wittmann, 2003; Slotkin et al., 2012). Understanding these chromosomal evolutionary processes provides essential context for grass systematics, ecology, and crop improvement, highlighting the continued relevance of cytogenetic research in modern plant biology (Grant, 1981; Raven, 1975).

## Acknowledgments

We thank CAPES and FAPEMIG for financial support. This study was conducted as part of the Plant Biology Graduate Program at Universidade Federal de Uberlândia.

## Author Contributions

ATHB compiled chromosomal data, performed analyses, and drafted the manuscript; CADW conceived the study, supervised the research, and revised the manuscript; CS assisted with analyses and manuscript revision.

